# UBA1 Mutations Drive RIPK1-Mediated Cell Death and Monocyte Dysfunction in VEXAS Syndrome

**DOI:** 10.1101/2025.10.06.680650

**Authors:** Paul Breillat, Samuel J. Magaziner, Stéphane Camus, Léa Dionet, Benjamin de Valence, Pierre Sohier, Amine Majdi, Quentin Delcros, Federica Pallotti, Nadia Rivet, Kevin Chevalier, Margot Poux, Pierre-Louis Tharaux, Olivia Lenoir, Abdelrahim Zoued, Olivier Kosmider, David B. Beck, Benjamin Terrier

## Abstract

VEXAS syndrome is a severe adult-onset autoinflammatory disease caused by somatic mutations in *UBA1* gene, disrupting cytoplasmic ubiquitin-activating enzyme E1 function in hematopoietic progenitors. The pathogenesis remains poorly understood, particularly how *UBA1* mutations perturb myeloid function. Here, we combine a genetically engineered THP-1 monocytic model with *ex vivo* analyses of blood and tissue samples from VEXAS patients to investigate the consequences of the canonical *UBA1*^M41V^ mutation. We show that *UBA1*-mutated monocytes exhibit TNF-α-induced cell death, characterized by RIPK1 phosphorylation, and MLKL-and caspase-8–mediated cell death. This is associated with defective transcriptional induction of NF-κB target genes and reduced cFLIP(L) expression in response to TNF-α. Monocytes also display blunted cytokine responses to multiple Toll-like receptor (TLR) agonists despite preserved TLR expression, linked to an impaired NF-κB response. UBA1^M41V^-derived macrophages exhibit an inflammatory transcriptional profile and increased secretion of chemokines that promote monocyte recruitment. We demonstrate that these UBA1^M41V^ macrophages display impaired efferocytosis due to lysosomal dysfunction. Together, these findings reveal a pathogenic axis in VEXAS syndrome linking UBA1 loss of function and defective ubiquitination to RIPK1-mediated inflammatory cell death, impaired antimicrobial signaling, and defective resolution mechanisms. Our study provides novel mechanistic insights into the myeloid dysfunction that drives inflammation and cytopenia in VEXAS and highlights the necroptosis and efferocytosis pathways as potential therapeutic targets.

**Key points:** 1/ *UBA1*-mutated monocytes are susceptible to RIPK1 dependent cell death and display impaired NF-κB–mediated cytokine responses to TLR agonists.

2/ UBA1-mutated macrophages promote inflammation and chemokine-mediated monocyte recruitment, while exhibiting defective efferocytosis.

## Introduction

VEXAS syndrome (Vacuoles, E1 enzyme, X-linked, Autoinflammatory, Somatic) is a recently characterized adult-onset autoinflammatory disease caused by somatic point mutations in the *UBA1* gene encoding the most abundant E1 ubiquitin ligase expressed in humans ^1^. These mutations predominantly affect methionine 41 (p.Met41) and lead to the expression of a truncated, nonfunctional UBA1c isoform, resulting in defects in ubiquitination ^1–3^. VEXAS syndrome is now recognized as a disorder affecting approximately 1 in 4,000 men over the age of 50 ^4^. It is characterized by systemic inflammation and hematologic complications, including cytopenias and myelodysplastic syndromes ^5–7^. Additionally, there is a high incidence of opportunistic infections, which contribute to disease-associated mortality ^8–11^.

The pathophysiology of VEXAS remains poorly understood. *UBA1* mutations are restricted to hematopoietic stem and progenitor cells (HSPCs) and persist in mature circulating monocytes ^1^ and myeloid cells that infiltrate inflamed tissues ^12,13^. While monogenic autoinflammatory diseases are typically caused by germline mutations in regulators of innate immunity, VEXAS represents a paradigm-shifting model where acquired defects in post-translational protein modification, drive dysregulated inflammation, cytopenias, and susceptibility to infections. Ubiquitination governs fundamental cellular processes, including protein degradation, cytokine signaling, and programmed cell death. These processes are critical to the function of myeloid cells and host defense. It is also essential for regulating the inflammatory response^14,15^.

Recent studies, including ours, have shown that circulating monocytes from VEXAS patients display both quantitative and qualitative defects, including altered inflammatory gene expression, features of functional exhaustion, and signatures suggestive of regulated cell death ^16–18^. However, mechanistic insights into how *UBA1* mutations perturb monocyte and macrophage function remain scarce. Prior investigations have primarily relied on single-cell transcriptomics of patient blood or bone marrow cells, and only rarely functional analyses on cellular models ^18–22^.

In this study, we aimed to delineate the functional consequences of the VEXAS-associated *UBA1^M41V^* mutation on human monocytes and macrophages. We used a genetically engineered THP-1 monocytic cell line model with conditional *UBA1* knock-out and stable expression of either wild-type or M41V-mutated *UBA1*. By combining this system with *ex vivo* analyses of blood and tissue samples from VEXAS patients, we uncover how the *UBA1^M41V^* mutation alters inflammatory responses, TNF-α-mediated cell death, and efferocytosis function in myeloid cells. Our findings reveal a pathogenic axis linking dysregulated ubiquitination, inflammatory cell death, and impaired macrophage host defense. This provides novel mechanistic insights into VEXAS syndrome and identifies potential therapeutic targets.

## Results

### Inflammatory cell death pathways are activated in myeloid cells from VEXAS patients

VEXAS syndrome is characterized by marked monocytopenia **(Figure 1A, Supplemental Table 1)** and distinctive cutaneous lesions that resemble histiocytoid Sweet’s syndrome ^23^. These lesions display a prominent infiltrate of CD68□ myeloid cells (**Supplemental Figure 1**), a hallmark of the disease’s inflammatory phenotype ^12,13^. Histological examination of VEXAS skin lesions revealed numerous karyorrhectic nuclei and extensive apoptotic debris underscoring an exacerbated local cell death process (**Figure 1B).**

**Figure. 1.**
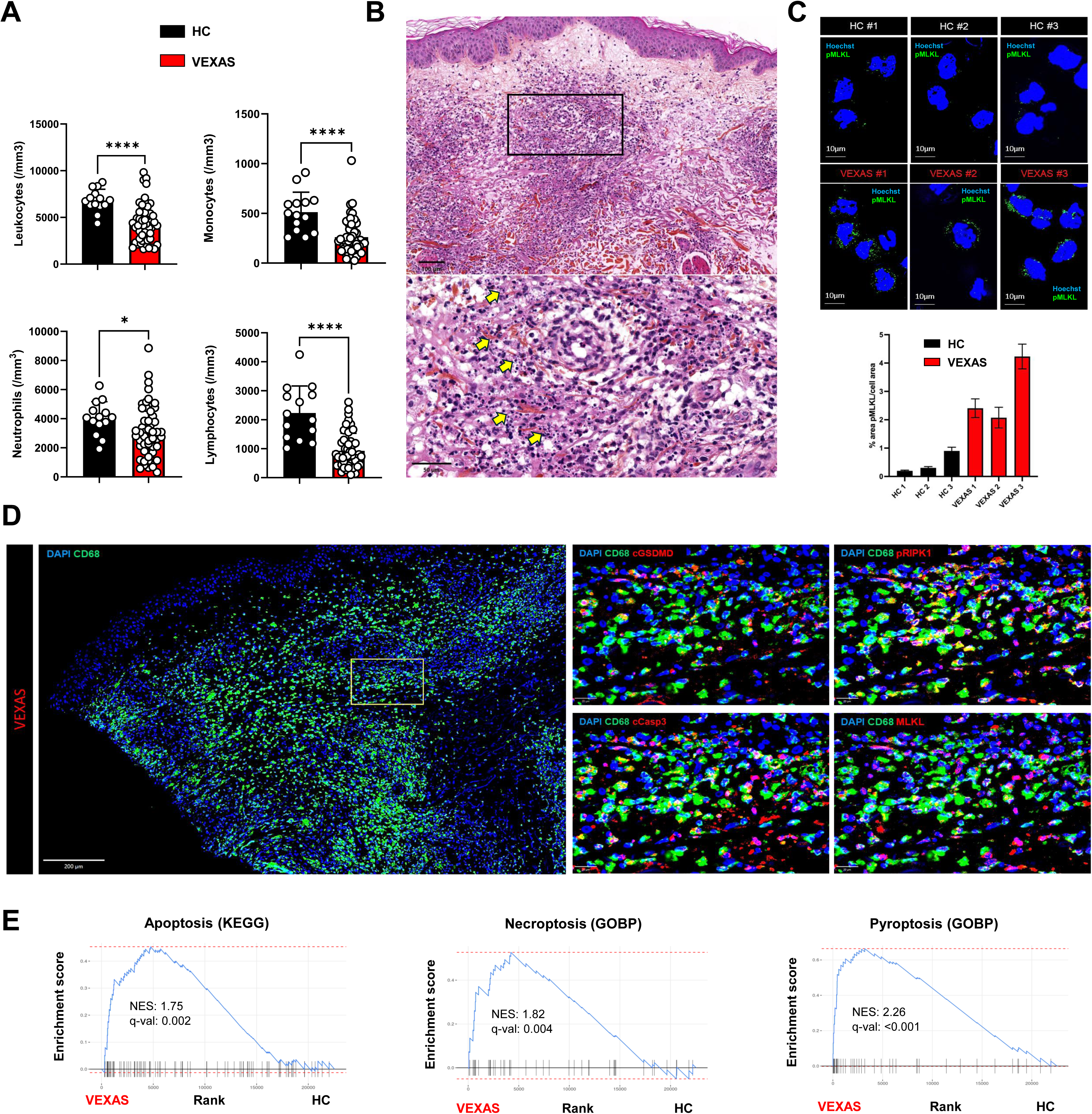
Regulated cell death pathways are activated in myeloid cells from VEXAS patients. (**A**) Leucocytes, neutrophils, monocytes, lymphocytes count from patients with VEXAS and elderly gender-matched healthy controls (HC). Panels show individual data (dots) and means ± SEM (histograms). P values were determined by the Mann-Whitney test. (**B**) Hematoxylin-eosin staining of UBA1-mutated Sweet-like lesion revealing karyorrhectic nuclei and apoptotic debris in VEXAS (arrows). Illustrative picture is shown (Magnification x10 and x40, scale bar 100 μm and 50µm). (**C**) Representative immunofluorescence images of pMLKL (green) staining on CD14+ sorted cells form 3 active VEXAS patients and 3 elderly gender-matched HC. Nuclei are stained in blue with Hoechst. Percentage of pMLKL cell surface area per CD14+ cells, data are shown and means (Histograms) ± SEM. Forty cells were quantified for each patient. (**D**) Multiplex Immunofluorescence of skin biopsy samples from VEXAS skin lesion showing the co-expression of cleaved GSDMD, MLKL, cleaved caspase-3 and phosphorylated RIPK1 within CD68+ infiltrates in VEXAS. (**E**) Gene set enrichment analysis of apoptosis (KEGG), necroptosis (GOBP) and pyroptosis (GOBP) pathways enriched in VEXAS lesional skin (n=6) versus non lesional skin from HC (n=5) adapted form from dataset GSE245639. *P□<□0.05; **P□<□0.01; ***P□<□0.001, ****P□<□0.0001. SEM, HC, Healthy Control; Standard Error of the Mean; VEXAS, Vacuoles, E1 enzyme, X-linked, Autoinflammatory, Somatic; WT, Wild-Type.

To characterize these mechanisms, we first evaluated the activation of programmed cell death pathways in peripheral myeloid cells. Single-cell RNA sequencing (scRNA-seq) of circulating monocytes revealed the upregulation of inflammatory signaling genes and multiple regulated cell death signatures, including necroptosis, apoptosis and pyroptosis ^16^. Necroptosis is a form of lytic, pro-inflammatory cell death initiated by TNF-α and mediated by RIPK1 and MLKL, linked with chronic inflammatory disorders ^24^. To investigate this pathway, we evaluated the phosphorylation of MLKL, the terminal effector of necroptosis, in CD14□ monocytes sorted from the peripheral blood of VEXAS patients. We observed increased levels of phosphorylated MLKL (pMLKL) in these cells, consistent with necroptosis activation *in vivo* (**Figure 1C**).

To confirm these findings at the tissue level, we performed multiplex immunofluorescence staining on skin biopsies from patients with VEXAS-histiocytoid Sweet’s syndrome. We observed extensive overlap of MLKL, cleaved caspase-3, phosphorylated RIPK1 and cleaved gasdermin D (GSDMD) within CD68□ infiltrates, supporting the activation of cell death initiated by TNF-α and mediated by RIPK1 in tissue-infiltrating myeloid cells (**Figure 1D**). In addition, transcriptomic profiling of these lesional biopsies from VEXAS patients confirmed upregulation of regulated cell death programs, including necroptosis, apoptosis and pyroptosis (**Figure 1E).** These results highlight a coordinated activation of inflammatory death pathways in both circulating and tissue-resident myeloid cells in VEXAS syndrome.

### UBA1 inactivation sensitizes monocytes to TNF-**α**-induced cell death

TNF-α signaling plays a dual role in host defense and autoinflammatory diseases, balancing survival through NF-κB activation with programmed cell death via apoptotic and necroptotic pathways, finely regulated by ubiquitination ^24–26^. Given that VEXAS syndrome is caused by somatic mutations in the ubiquitin-activating enzyme UBA1, which orchestrates the initiation of most ubiquitin-dependent signaling cascades, we sought to examine the specific contribution of UBA1 inactivation, and in particular the canonical UBA1^M41V^ mutation, to inflammatory cell death in monocytes.

To this end, we used a THP-1 monocytic model with a doxycycline-inducible knockout of endogenous UBA1 (UBA1^KO^), completed with constitutive expression of the UBA1^WT^ or UBA1^M41V^ variant. Following induction with doxycycline, UBA1^KO^ cells exhibited complete loss of both nuclear and cytoplasmic UBA1 isoforms (UBA1a and UBA1b), leading to a global reduction in polyubiquitin chain formation (**Supplementary** Figure 2A). In parallel, UBA1^M41V^ THP-1 cells expressed the truncated cytoplasmic UBA1c isoform, lacked functional UBA1b, and displayed defective polyubiquitination (**Supplementary** Figure 2B). Morphological assessment revealed cytoplasmic vacuolization in UBA1^M41V^ cells, recapitulating one of the hallmarks of VEXAS syndrome **(Supplementary** Figure 2C) Transcriptomic profiling of UBA1^M41V^ THP-1 cells showed enrichment of unfolded protein response, pro-inflammatory response pathways, and regulated cell death programs, including necroptosis and apoptosis, relative to UBA1^WT^ THP-1 cells (**Figure 2A and 2B**). Consistent with these findings, spontaneous cell death progressively increased over time in UBA1^KO^ and, to a lesser extent, UBA1^M41V^ cells after induction (**Figure 2C**). Upon TNF-α stimulation, both UBA1^KO^ and UBA1^M41V^ THP-1 cells showed increased susceptibility to cell death, as evidenced by increased propidium iodide (PI) incorporation and Annexin V staining compared to UBA1^WT^ cells (**Figure 2D and 2E**). This suggests a cell-intrinsic vulnerability to TNF-α-induced cell death associated with UBA1 inactivation.

**Figure. 2.**
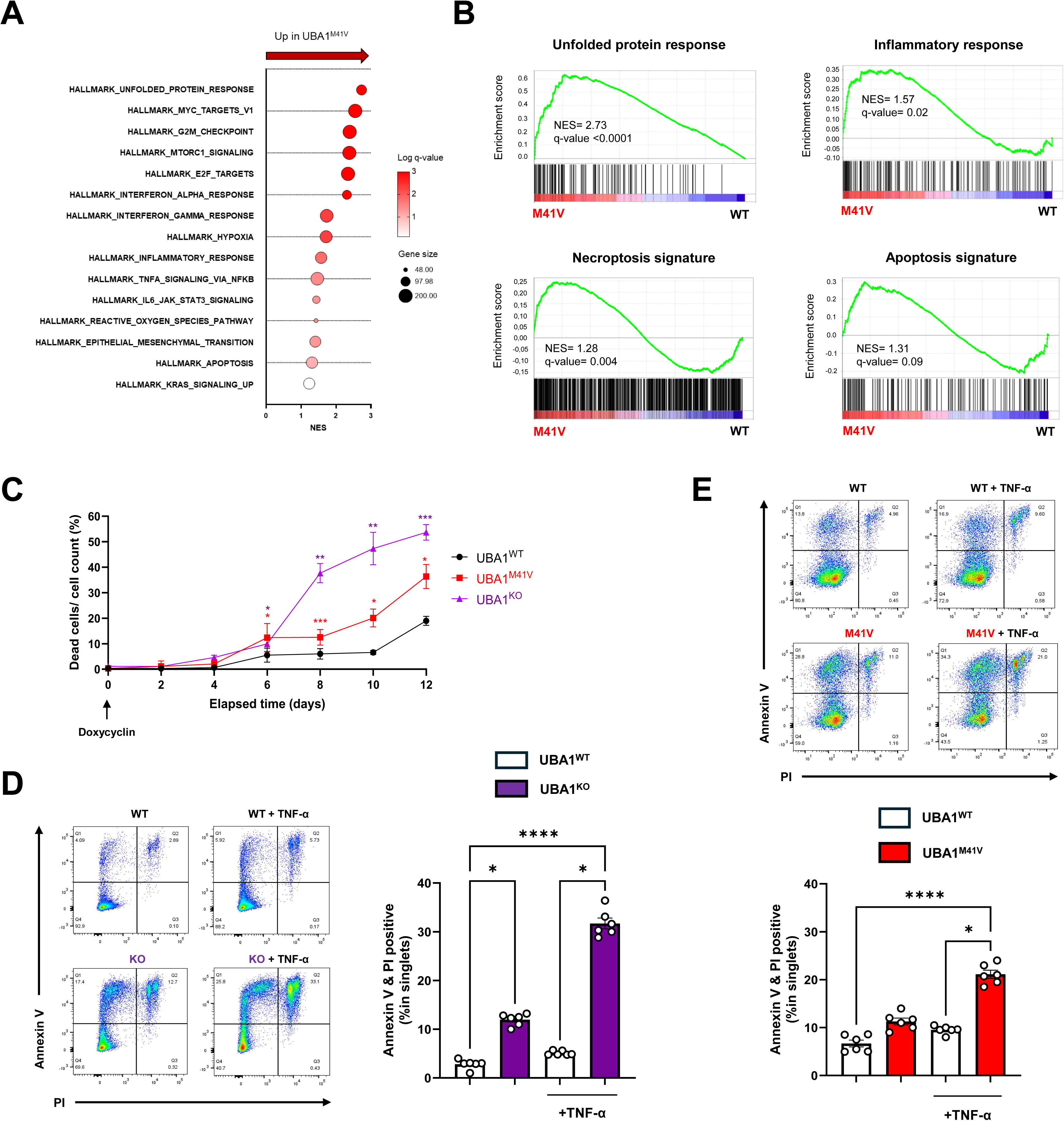
UBA1 inactivation sensitizes monocytes to TNF**α**-induced cell death. (**A**) Representation of 15 upregulated Hallmark pathways in UBA1^M41V^ compared to UBA1^WT^ THP-1 cells (n = 4 independently collected samples). Log (10) of q-value=0 was arbitrary set at 3. (**B**) GSEA analysis of Unfold Protein Response (Hallmark), Inflammatory Response (Hallmark), Apoptosis (Hallmark) and Necroptosis (Culver-Cochran et al, Nature Communications, 2024) from in UBA1^WT^ and UBA1^M41V^ THP-1 cells. (**C**) Time-lapse quantification of dead cell percentages in UBA1^WT^, UBA1^KO^ and UBA1^M41V^ cells after doxycycline induction (Data represent at least 3 independent experiments) using trypan blue. The viabilities in UBA1^M41V^ and UBA1^KO^ cells were compared to the viability of UBA1^WT^ cells at each time point with unpaired t-tests. (**D**) Representative flow plots of annexin V and PI detection by Flow Cytometry in UBA1^WT^ and UBA1^KO^ THP-1 cells. (**E**) Representative flow plots of annexin V and PI detection by Flow Cytometry in UBA1^WT^ and UBA1^M41V^ THP-1 cells. THP-1 cells were stimulated with TNFα 24 hours or left unstimulated. 2 tailed p-values were determined with non-parametric Kruskal-Wallis test, followed by Dunn’s post test for multiple group comparisons. *P□<□0.05; **P□<□0.01; ***P□<□0.001, ****P□<□0.0001 SEM, Standard Error of the Mean; WT, wild-type;

To investigate whether loss of UBA1 function triggers regulated inflammatory cell death, we used a human monocytic reporter cell line (THP1-HMGB1-Lucia) engineered to detect extracellular release of HMGB1, a prototypical damage-associated molecular pattern (DAMP) associated with pore-forming forms of cell death such as necroptosis ^25^. Treatment with the selective E1 enzyme inhibitor PYR-41, in combination with TNF-α, induced HMGB1 release, consistent with the induction of lytic cell death (**Figure 3A**). Importantly, this effect was almost completely abrogated by co-treatment with the RIPK1 inhibitor necrostatin-1 or the MLKL inhibitor necrosulfonamide, suggesting RIPK1-and MLKL-dependent cell death as the dominant pathway in this context (**Figure 3A**). Next, we assessed these signaling events in our *UBA1*-deficient cell models. Western blot analysis of UBA1^KO^ and UBA1^M41V^ THP-1 cells stimulated with TNF-α for 24 hours revealed increased phosphorylation of RIPK1, as well as cleavage of caspase-8 and caspase-3 in treated cells compared to UBA1^WT^ (**Figure 3B and 3C**), confirming the engagement of apoptosis downstream of TNFR1. Consistent with activation of necroptosis, immunofluorescence analysis showed phosphorylation and membrane translocation of MLKL in UBA1^M41V^ THP-1 cells after 3 hours of TNF-α exposure. This process was inhibited by necrostatin-1 pre-exposure (**Figure 3D**). Quantification of cell death following TNF-α stimulation, either with or without inhibition of inhibitors of cIAP and XIAP (BV-6) and pan-caspases inhibitor (Z-VAD-FMK), confirms that UBA1^M41V^ THP-1 cells are sensitized to TNF-α-dependent cell death, particularly necroptosis. Furthermore, TNF-α–induced cell death in UBA1^M41V^ cells was only partially reduced by pan-caspase inhibition (Z-VAD-FMK) and almost completely blocked by RIPK1 inhibition (necrostatin-1), MLKL inhibition (necrosulfonamide), or dual caspase and MLKL inhibition (**Figure 3E**). We further explored the functional consequences of this inflammatory cell death by quantifying cytokine secretion in the supernatant of TNF-α-stimulated THP-1 cells. Compared to UBA1^WT^ controls, both UBA1^KO^ and UBA1^M41V^ THP-1 cells secreted significantly higher levels of IL-1β and IL-8 inflammatory mediators. Notably, pretreatment with necrostatin-1 markedly reduced this cytokine release (**Figure 3F**), supporting the role of RIPK1 kinase activity in promoting both cell death and inflammatory signaling.

**Figure. 3.**
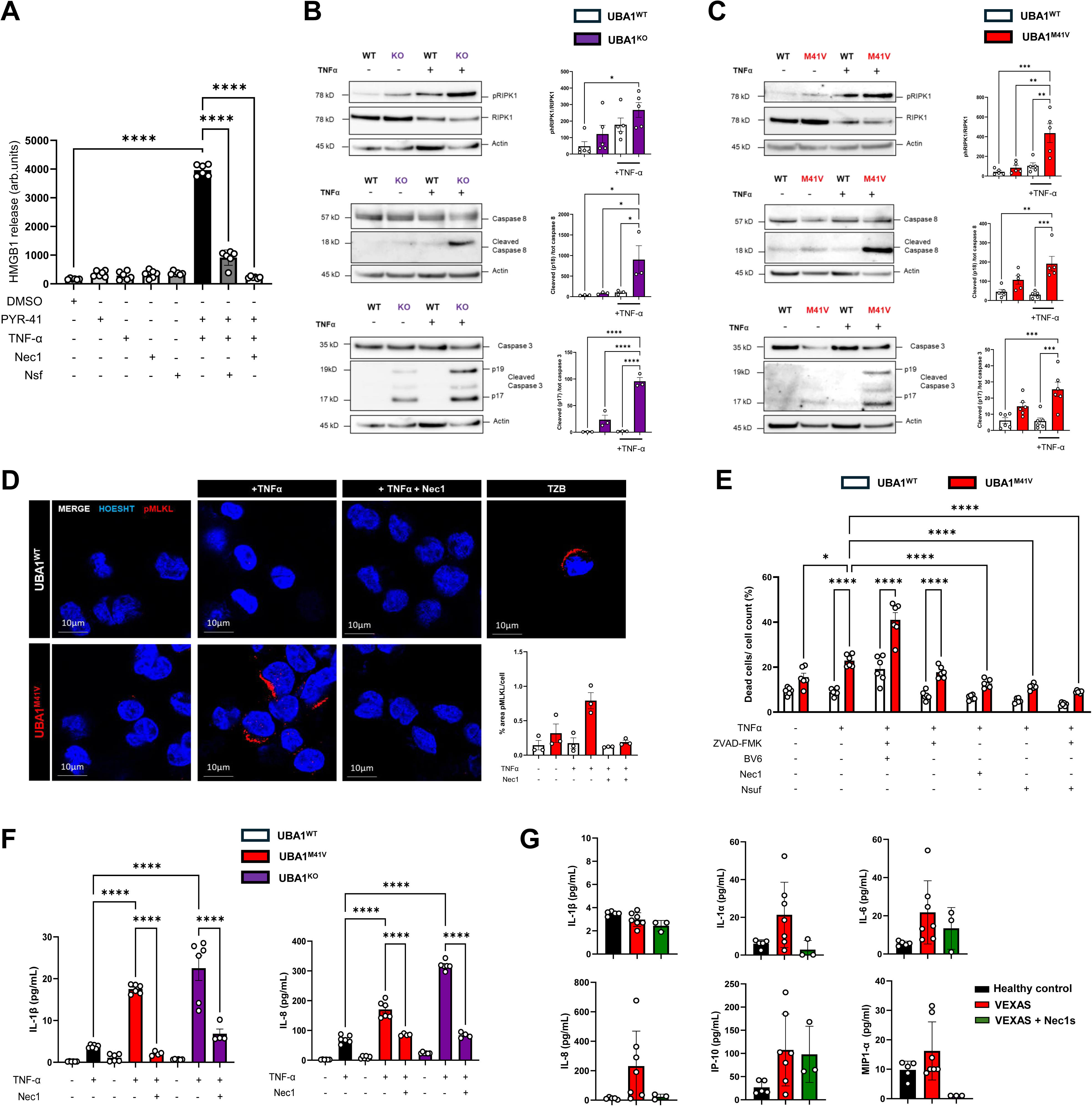
TNFα-induced cell death is mediated by RIPK1 and drives inflammation in *UBA1*-mutated cells. (**A**) THP-1-HMGB1-Lucia cells were pre-treated with a selective inhibitor of ubiquitin-activating enzyme E1 PYR-41 (5µM) with or without Necrostatin-1 (20µM) or Necrosulfamide (1µM) for 1 hour prior to incubation with TNF-α (50 ng/ml). Twenty-four hours later, the luciferase activity was determined by measuring the luminescence in a luminometer. Analysis was performed using non-parametric ANOVA test followed by Dunn’s post-test, (Not all P values are represented). (**B and C**) TNF-α-induced RIPK1 phosphorylation and caspase 8 and 3 cleavages assessed by immunoblotting in UBA1^WT^, UBA1^KO^ (**B**) and UBA1^M41V^ (**C**) THP-1 cells. THP-1 cells were stimulated with TNF-α for 24 hours or left unstimulated. Analysis was performed using non-parametric ANOVA test followed by Dunn’s post-test (at least *n*□=□3 independent experiment). (**D**) Representative immunofluorescence images of pMLKL (red) staining on THP-1 UBA1^WT^ and UBA1^M41V^ cells. Nuclei are stained in blue with Hoechst. THP-1 cells were stimulated 3h with TNF-α (T), BV-6 (B), Z-VAD-FMK (Z), Necrostatin-1 (Nec1). TZB condition served as a positive control for necroptosis. Unstimulated controls were treated with DMSO. Percentage of pMLKL cell surface area per THP1 cells was calculated and each dot represents the mean of 40 cells quantification in 3 independents experiments. Panels show means ± SEM. (**E**) UBA1^WT^ and UBA1^M41V^ THP-1 cells were stimulated with TNFα (50ng/ml), BV-6 (5µM), Z-VAD-FMK (20µM), Nec1 (20µM) and Necrosulfonamide (Nsuf) (1µM) for 24 hours and the percentage of living cells was determined by Zombie staining assessed with the Celigo Image Cytometer. Unstimulated controls were treated with DMSO. Each condition was performed in triplicate. Data represent the results of 6 independent experiments. Analysis was performed using non-parametric ANOVA test followed by Dunn’s post-test, (Not all P values are represented). (**F**) IL-1β and IL-8 levels in supernatant from UBA1^KO^ and UBA1^M41V^ THP-1 cells after 24H TNF-α (50ng/ml) stimulation compared to UBA1^WT^ controls in presence or not of Nec1(20µM). Unstimulated controls were treated with DMSO. (**G)** VEXAS patients exhibited elevated secretion of IL-1α, IL-8, and MIP-1α, which was attenuated by addition of Necrostatin-1S (Nec1S). Whole blood from VEXAS patients and HCs was incubated 22 hours in TruCulture tubes preloaded with Nec1S (10µM) or unstimulated. Supernatant were harvested, and cytokine levels determined using Luminex. *P□<□0.05; **P□<□0.01; ***P□<□0.001, ****P□<□0.0001 SEM, Standard Error of the Mean; WT, wild-type; Necrostatin-1, Nec1; Necrostatin-1S, Nec1S; Necrosulfonamide, Nsuf.

Since recent findings indicate that TNFR1 signaling can lead to pyroptotic cell death via the caspase-8-mediated cleavage of GSDMD in a RIPK1-dependent manner ^27,28^, we examined GSDMD processing in our model. Following TNF-α stimulation, we detected GSDMD cleavage in UBA1^KO^ and UBA1^M41V^ THP-1 cells, though at a lower level in UBA1^M41V^ than in UBA1^WT^ THP-1 cells (**Supplementary** Figure 3). These results suggest that pyroptosis is not a major contributor to cell death in this context, but rather a potential additional contributor to cell death and inflammation.

To validate these findings in a patient-derived context, we performed whole-blood stimulation assays using the standardized TruCulture platform, a standardized *ex vivo* whole-blood assay that preserves innate immune cell interactions (**Supplementary Table 2 for main characteristics of patients**). After 22 hours of culture, blood samples from VEXAS patients exhibited elevated secretion of IL-1α, IL-8, and MIP-1α, which was attenuated by addition of necrostatin-1s (**Figure 3G**). These results support a model in which *UBA1*-mutated monocytes undergo RIPK1-dependent necroptosis in response to TNF-α, thereby contributing to systemic inflammation.

### UBA1^M41V^ mutation impairs NF-**κ**B-dependent signaling and cFLIP expression downstream of TNF-**α**

To better understand the increased susceptibility of *UBA1*-mutant monocytes to TNF-α-induced cell death, we examined the integrity of the canonical NF-κB signaling pathway, a major survival axis downstream of TNFR1 activation. In response to TNF-α stimulation, UBA1^M41V^ THP-1 cells exhibited preserved phosphorylation of the p65 subunit but delayed degradation of IκBα, the cytoplasmic inhibitor of NF-κB (**Figure 4A**), potentially impairing nuclear translocation and transcriptional activation of NF-κB target genes. To validate this observation functionally, we employed a monocytic NF-κB reporter cell line (THP1-NF-κB-Blue), which demonstrated that inhibition of E1 enzyme activity by PYR-41 caused a dose-dependent suppression of NF-κB-driven transcription in response to TNF-α (**Figure 4B**). This effect persisted even in the presence of caspase and RIPK1 inhibitors (Z-VAD-FMK and necrostatin-1, respectively), indicating that the transcriptional defect occurs upstream of complex II-mediated cell death (**Figure 4B**). Consistent with these findings, UBA1^M41V^ THP-1 cells displayed impaired induction of classical NF-κB target genes (*TNFAIP3, ICAM1, NFKBIA, BIRC3, TNFAIP6*) following TNF-α stimulation, as evidenced by bulk RNA-seq analysis (**Figure 4C and 4D**).

**Figure. 4.**
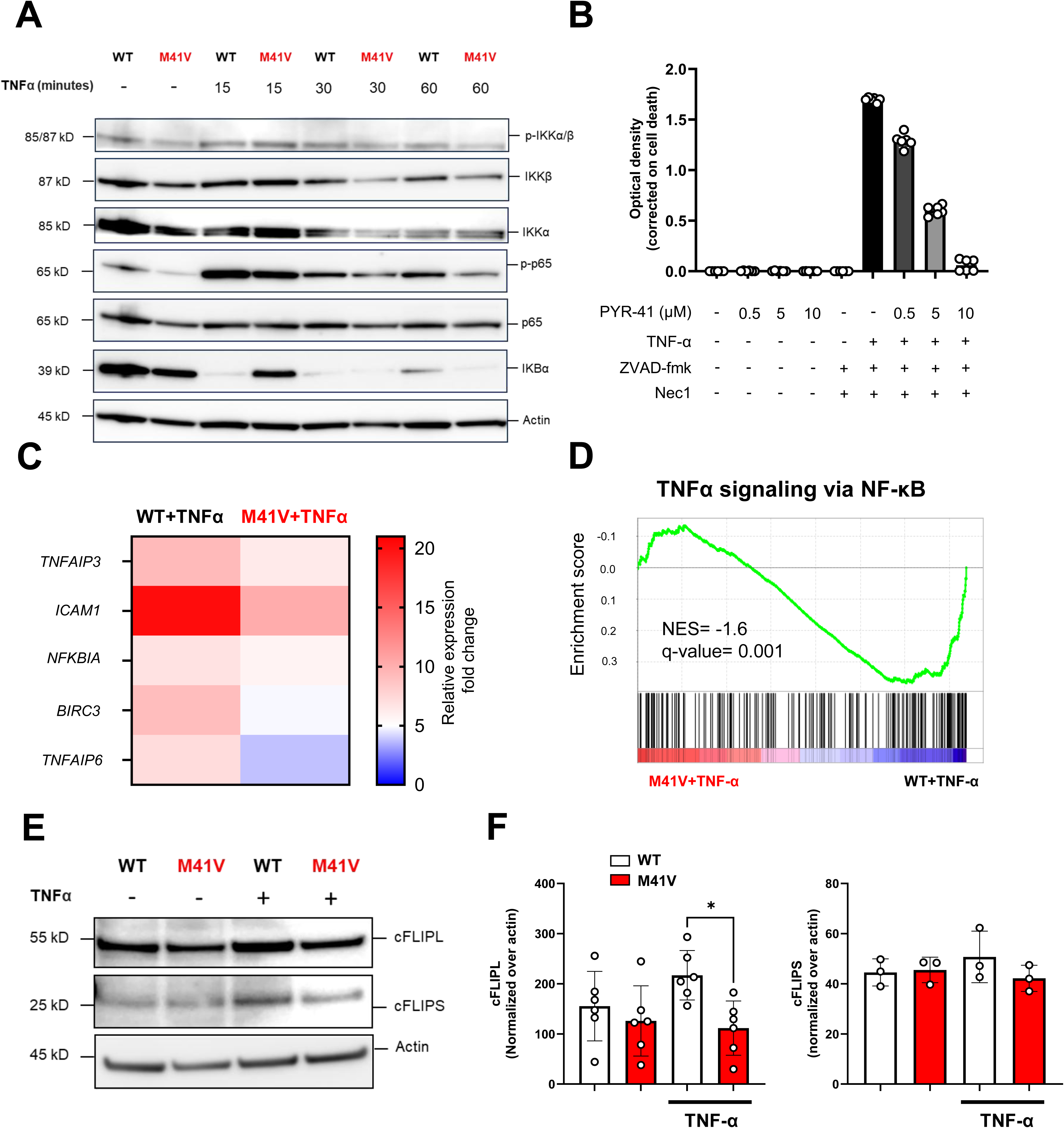
Impaired NF-**κ**B signal transduction and low cFLIP(L) expression in UBA1^M41V^ THP-1 cells after TNF-**α** challenge. **(A**) Immunoblot analysis of NF-κB pathway in UBA1^WT^ and UBA1^M41V^ THP-1 cells. Cells were challenge with TNFα (50mg/ml) 15, 30 and 60 min or left untreated. One blot representative of 3 independent experiments is shown. (**B**) NF-κB signal transduction after TNFα (50mg/ml) 24H challenge in a context of ubiquitin-activating enzyme E1 inhibition with PYR-41(5µM) using NF-κB reporter THP-1 cell line (THP-1-Blue-NF-κB, Invivogen). Unstimulated controls were treated with DMSO. Cells were pretreated with Nec1 (20µM) and Z-VAD-FMK (20µM) to prevent cell death and data were normalized on cell viability (*n*□=□6 independent experiments). (**C**) UBA1^WT^ and UBA1^M41V^ THP-1 cells were treated with TNF-α (50□ng/ml) for 24H or let unstimulated, mRNA transcription change of NF-κB targeted genes were determined and compared to UBA1^WT^ non stimulated cells as reference (*n=*4 independent experiments). (**D**) GSEA analysis of TNF-α signaling via NF-κB (Hallmark) in UBA1^WT^ and UBA1^M41V^ THP-1 cells after TNF-α (50□ng/ml) challenge for 24H (n=4 independent experiments). (**E**) Representative immunoblot of cFLIP isoforms in UBA1^WT^ and UBA1^M41V^ THP-1 cells. Cells were treated with TNF-α (50□ng/ml) for 24H or left unstimulated. (**F**) Quantification of cFLIP isoforms in UBA1^WT^ and UBA1^M41V^ THP-1 cells. At least n=3 independent experiments are represented and 2 tailed p-values were determined with non-parametric ANOVA test followed by Dunn’s post-test. *P□<□0.05; **P□<□0.01; ***P□<□0.001, ****P□<□0.0001 SEM, Standard Error of the Mean; WT, wild-type; Necrostatin-1, Nec1

Given the role of cFLIP as a key checkpoint regulating caspase-8 activity at the Death-Inducing Signaling Complex (DISC) and induced by NF-κB signaling ^29,30^, we analyzed cFLIP expression by immunoblot. We observed a marked reduction in the long isoform (cFLIP_L) of the protein in UBA1^M41V^ THP-1 cells upon TNF-α stimulation, while expression of the short isoform (cFLIP_S) remained unchanged (**Figure 4E and 4F**).

Collectively, these results support a model in which *UBA1* mutations impair NF-κB transcriptional activation downstream of TNF-α, in part through defective IκBα processing and reduced cFLIP_L expression, and promote cell-death through RIPK1-mediated cell death.

### Impaired TLR-mediated inflammatory responses in *UBA1*-mutated monocytes

Given the high incidence of opportunistic infections in VEXAS syndrome, we hypothesized that monocytes may exhibit defective responses to pathogen-associated molecular patterns (PAMPs), particularly via Toll-like receptors (TLRs), which are central to innate immune sensing and cytokine production ^31^. To address this, we performed whole-blood stimulation assays ^32^ using a panel of TLR agonists, poly(I:C) (TLR3), lipopolysaccharide (LPS; TLR4), and R848 (TLR7/8), in patients with genetically confirmed VEXAS syndrome (**Supplementary Table 2**).

After 22 hours of stimulation, blood samples from VEXAS patients showed significantly lower production of key pro-inflammatory cytokines. Compared to controls, secretion of IL-1β, IL-6, IFN-γ, and TNF-α was markedly decreased in response to LPS or Poly(I:C) stimulation, while R848 stimulation elicited a trend toward reduced IL-6 and IFN-γ secretion though the response was more variable (**Figure 5A**). These results suggest that VEXAS syndrome could be associated with a broad impairment of TLR-mediated innate responses.

**Figure. 5.**
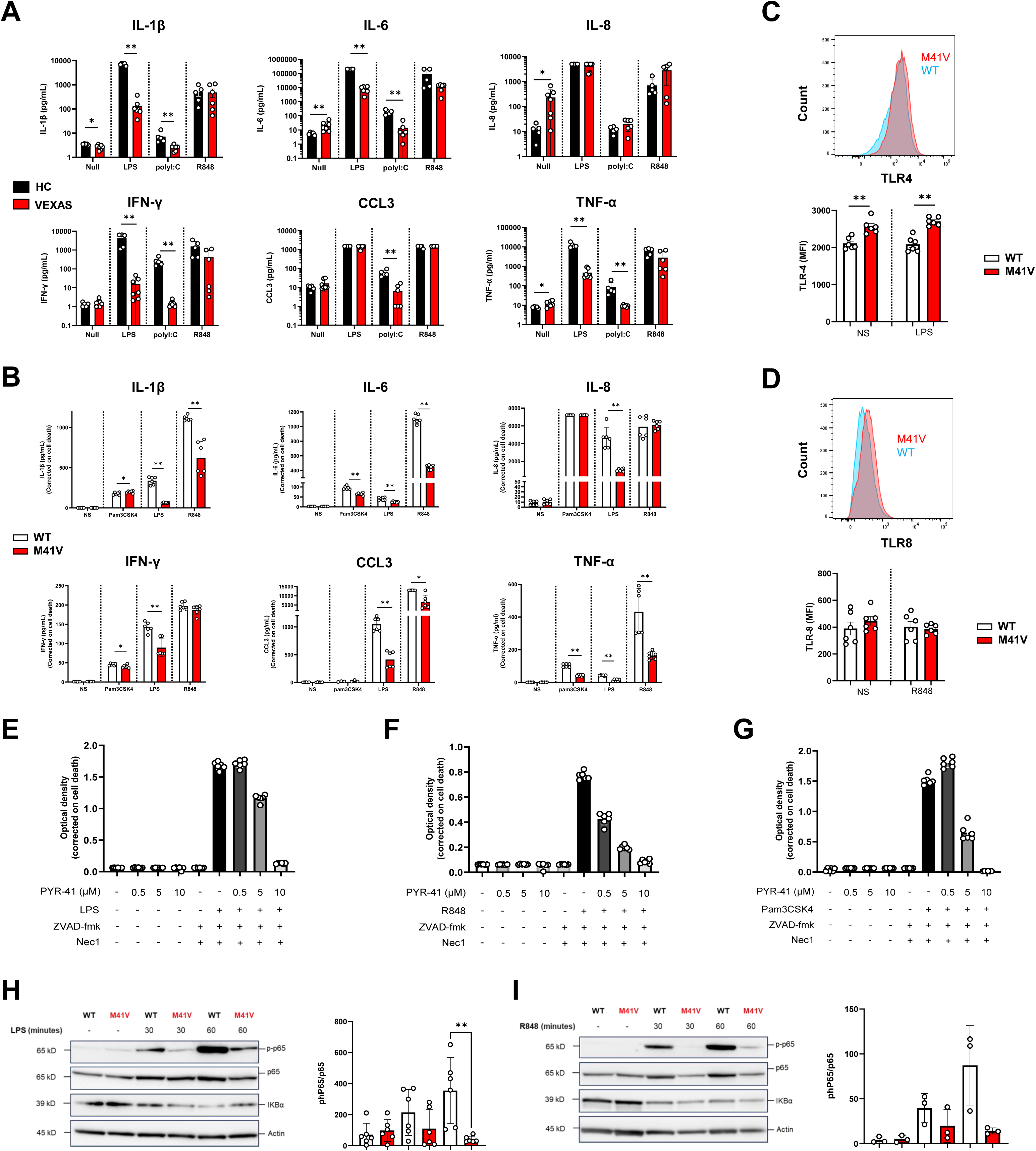
Impaired TLR-mediated inflammatory responses in UBA1^M41V^ monocytes. **(A**) Cytokine and chemokines levels in whole blood culture supernatants from VEXAS and healthy controls at baseline (Null) and after 22H challenge with TLR-3 agonist (poly(I:C), TLR-4 agonist (LPS), TLR-7/8 agonist (R848). Panels show individual data (dots) and means ± SEM (histograms). (**B**) Cytokine and chemokines levels in UBA1^WT^ and UBA1^M41V^ THP-1 cells after 24H challenge with TLR2 agonist (Pam3CSK4, 100ng/ml), TLR-4 agonist (LPS, 100ng/ml), TLR-7/8 agonist (R848, 100µg/ml). Data were normalized on cell viability. (**C**) TLR-4 extracellular expression in UBA1^WT^ and UBA1^M41V^ THP-1 cells assessed by flow cytometry. Cells were challenge with LPS (100ng/mL) 6H or left unstimulated. (**D**) TLR-8 intracellular staining in THP-1 cells assessed by flow cytometry. Cells were challenge with R848 (2.5µg/mL) 6H or left unstimulated. For (**C**) and (**D**), 6 independent experiments are represented and 2 tailed p-values were determined with Mann Witney test for Mean Fluorescence Intensity (MFI) comparision. (**E, F, G**) NF-κB signal transduction after TLR agonists (TLR2 (Pam3CSK4, 100ng/ml), TLR-4 (LPS, 100ng/ml), TLR-7/8 (R848, 100µg/ml)) challenge in a context of ubiquitin-activating enzyme E1 inhibition with PYR-41(5µM) using NF-κB reporter THP-1 cell line (THP-1-Blue-NF-κB, Invivogen). Unstimulated controls were treated with DMSO. Cells were pretreated with Nec1 (20µM) and Z-VAD-FMK (20µM) to prevent cell death and data were normalized on cell viability (*n*□=□6/group). (**H** and **I**) Study of NF-κB signaling pathway assessed by p65 phosphorylation and IKBα expression following LPS (100ng/ml) (**H**) and R848 (100µg/ml) (**I**) challenge by immunoblotting UBA1^WT^ and UBA1^M41V^ THP-1 cells. THP-1 cells were stimulated for 30, 60 minutes or left unstimulated. For (**H**) and (**I**), 6 and 3 independent experiments are represented respectively and 2 tailed p-values were determined with Mann Witney unpaired test. LPS, Lipopolysaccharides; poly(I:C), Polyinosinic-polycytidylic acid; R848, Resquimod; SEM, Standard Error of the Mean; TLR, Toll-Like receptor; VEXAS, Vacuoles, E1 enzyme, X-linked, Autoinflammatory, Somatic. *P□<□0.05; **P□<□0.01; ***P□<□0.001, ****P□<□0.0001.

To confirm whether this defect is intrinsic to *UBA1*-mutated monocytes, we stimulated UBA1^M41V^ THP-1 cells with the same TLR agonists. UBA1^M41V^ cells exhibited diminished secretion of IL-1β, IL-6, IFN-γ, CCL3, and TNF-α following stimulation with LPS or Pam3CSK4, a TLR1/2 ligand (**Figure 5B**). R848 did elicit a clear differential response between UBA1^M41V^ and UBA1^WT^ THP-1, with decreased secretion of IL-1β, IL-6, CCL3 ad TNF-α. As expected, no response was observed upon poly(I:C) stimulation due to the absence of TLR3 expression in THP-1 cells (data not shown). These data are consistent with patient data, confirming global impairment of TLR response to agonists.

To exclude the possibility that these defects were due to reduced receptor expression, we assessed the surface levels of TLR4 and TLR7/8 using flow cytometry. We only observed a slight increase in TLR4 expression in UBA1^M41V^ cells (**Figure 5C**) and no significant differences for TLR8 compared to UBA1^WT^ cells (TLR7 was not expressed in either WT or M41V cells (data not shown)) (**Figure 5D**), supporting a post-receptor signaling defect in UBA1^M41V^ cells. We next evaluated TLR-dependent NF-κB pathway activation using the monocytic NF-κB reporter cell line (THP1-NF-κB-Blue). Pharmacologic inhibition of E1 enzyme activity with PYR-41 resulted in a dose-dependent suppression of NF-κB-driven luciferase activity in response to LPS, R848, and Pam3CSK4 (**Figure 5E, 5F, and 5G**), consistent with impaired signal transduction downstream of TLRs upon UBA1 inhibition. Finally, Western blot analysis confirmed defective phosphorylation of the NF-κB subunit p65 upon LPS and R848 stimulation in UBA1^M41V^ THP-1 cells, further supporting a defect in NF-κB pathway activation (**Figure 5H and 5I**).

Together, these data demonstrate that *UBA1* mutations impair monocyte responsiveness to multiple TLR agonists, likely through defective downstream signaling, including the NF-κB pathway, thereby compromising antimicrobial defense and contributing to infection susceptibility in VEXAS syndrome.

### UBA1^M41V^ THP-1-derived macrophages exhibit a pro-inflammatory phenotype with defective efferocytosis

VEXAS syndrome is marked by systemic and tissue inflammation, with myeloid cell infiltration and dead cell debris accumulation, particularly within skin lesions, suggesting a role for dysfunctional macrophages in driving local immune activation and insufficient resolution of inflammation. To explore the impact of *UBA1* mutation in differentiated macrophages, we generated THP-1-derived macrophages from UBA1^M41V^ and UBA1^WT^ cells. UBA1^M41V^ THP-1-derived macrophages lacked the functional cytoplasmic UBA1b isoform and expressed the truncated UBA1c isoform (**Supplementary** Figure 4A). Culture supernatants from UBA1^M41V^-derived macrophages revealed a marked pro-inflammatory profile, with significantly elevated secretion of IL-1β, IL-6, IFN-γ, TNF-α, and CXCL10 compared to UBA1^WT^-derived macrophages (**Supplementary** Figure 4B). Bulk transcriptomic analysis further confirmed enrichment of inflammatory gene signatures, including pathways associated with TNF-α/NF-κB, type I and II interferons, the unfolded protein response, necroptosis and apoptosis (**Supplementary** Figure 4C and 4D). In line with these findings, UBA1^M41V^ macrophages exhibited increased cell death (**Supplementary** Figure 4E and 4F).

Following classical polarization with LPS and IFN-γ, UBA1^M41V^-derived macrophages retained and amplified their inflammatory phenotype. This was evidenced by increased expression of *NOS2* and *CXCL10* and secretion of IL-1β and IL-6. Paradoxically, we detected a decreased expression of co-stimulatory molecules *CD80* and *CD86* (**Supplementary** Figure 4G). In contrast, we noticed a marked impairment of anti-inflammatory phenotypic adjustment of UBA1^M41V^ macrophages after alternative polarization with IL-4 and IL-13 with decreased and expression of *CD163*, *CD206, IL10,* and *MERTK* and associated with impaired secretion of IL-10 (**Supplementary** Figure 4H), suggesting a defective immunoregulatory response.

Given the prominent cutaneous infiltration of myeloid cells in VEXAS and the potential for cell-intrinsic chemokine production to shape tissue tropism, we next assessed the chemotactic profile of *UBA1*-mutated macrophages. Plasma from patients with VEXAS (n=40) and culture supernatants from UBA1^M41V^-derived macrophages both showed elevated levels of key chemokines, including CCL-2 and CCL-20, compared to healthy donors (n=10) and UBA1^WT^ cells, respectively (**Supplementary** Figure 5A and 5B**, Supplementary Table 3**). In functional migration assays, UBA1^M41V^ cells displayed increased migration in response to IL-8, CCL2, and CCL3 (**Supplementary** Figure 5C). Moreover, supernatants from UBA1^M41V^-derived macrophages induced significantly greater THP-1 UBA1^WT^ and UBA1^M41V^ cells migration than those from UBA1^WT^ macrophages (**Supplementary** Figure 5D), suggesting that *UBA1*-mutated macrophages may preferentially attract clonal myeloid cells into inflamed tissues, thereby reinforcing the inflammatory loop and contributing to clonal dominance in affected compartments.

Given the accumulation of dead cell debris in VEXAS tissue biopsies and the notorious vulnerability of VEXAS patients to intracellular pathogens and fungus infections, we assessed the phagocytic and efferocytosis capacity of UBA1^M41V^-derived macrophages. Although surface expression of MERTK, a key receptor for apoptotic cell recognition, was reduced in UBA1^M41V^-derived macrophages (**Figure 6A**), the initial engulfment of apoptotic bodies was preserved. However, we observed an impairment in the clearance rate of engulfed apoptotic cells (**Figure 6B, Supplementary** Figure 6A).

**Figure. 6.**
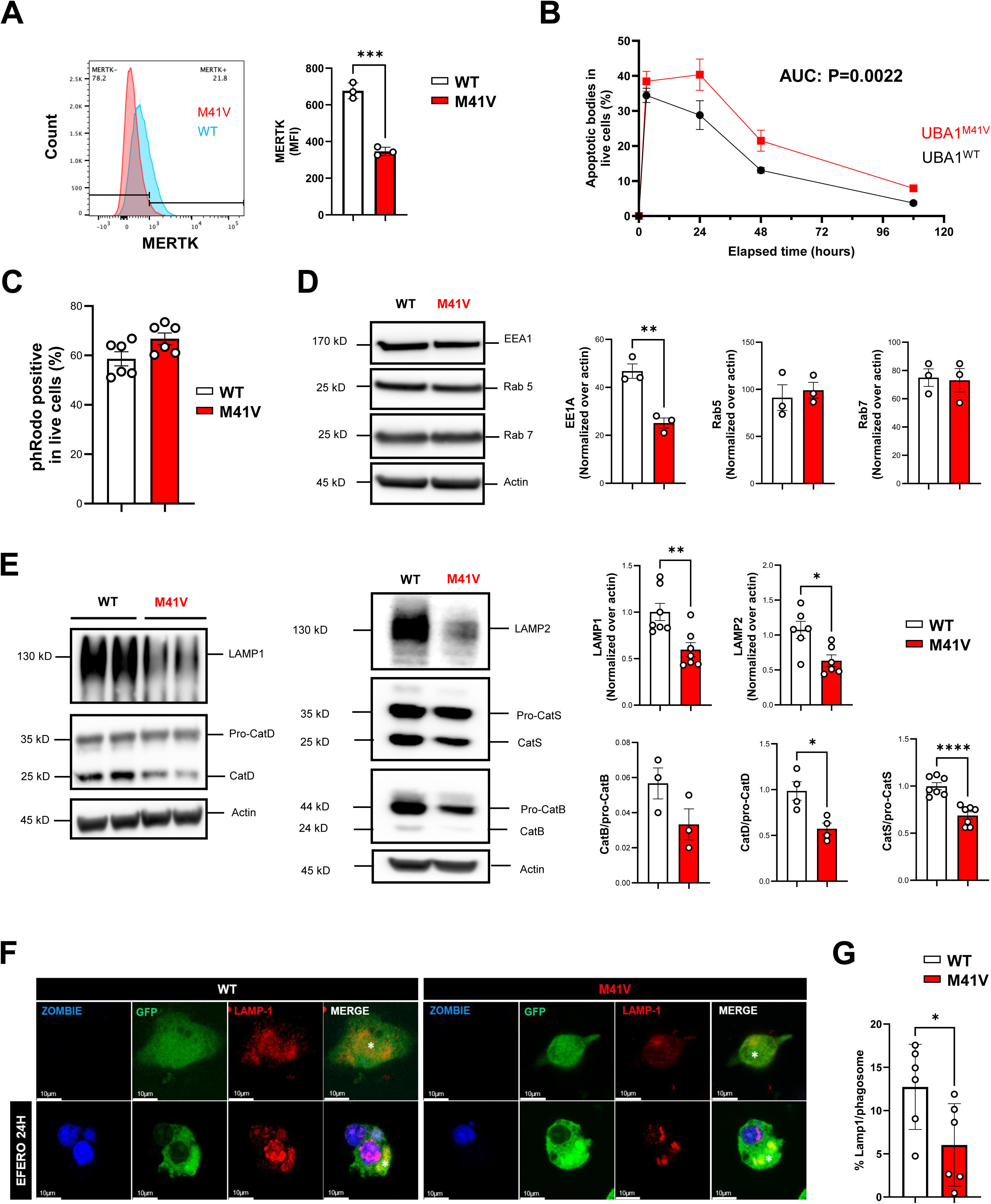
UBA1^M41V^ THP-1-derived macrophages exhibit impaired efferocytosis. (**A**) MERTK extracellular expression in UBA1^WT^ and UBA1^M41V^ THP-1 derived macrophages assessed by flow cytometry. Three independent experiments are represented and 2 tailed p-values were determined with unpaired t test for Mean Fluorescence Intensity (MFI) comparision. (**B**) Flow cytometric analysis of apoptotic bodies clearance in UBA1^WT^ and UBA1^M41V^ THP-1 derived macrophages using HeLa dead cells stained with zombie dye as target (1:1 ratio with live Macrophages). Data represent n=6 independent experiments at each time points. Area under the curve was calculated for each condition then compared using unpaired Mann Witney test. (**C**) Flow cytometric analysis of the phagocytic capacity of UBA1^WT^ and UBA1^M41V^ THP-1 derived macrophages using pHrodo *E. coli* as target particles. (**D**) Study of endocytosis effectors expression in UBA1^WT^ and UBA1^M41V^ THP-1 derived macrophages assessed by immunoblotting. Data represent 3 independent experiments and 2 tailed p-values were determined with Welch’s t test. (**E**) Study of late efferocytosis and lysosomal protease expression in UBA1^WT^ and UBA1^M41V^ THP-1 derived macrophages assessed by immunoblotting. Data represent at least 3 independent experiments and 2 tailed p-values were determined with Welch’s t test. (**F**) Representative immunofluorescence images of LAMP1staining (in red) in UBA1^WT^ and UBA1^M41V^ THP-1 derived macrophages (in green) co cultured 24H with HeLa dead cells stained with zombie dye (in blue) as target (1:1 ratio with live Macrophages). Asterix shows nuclear staining. (**G**) Quantification of LAMP1 % of phagosome surface in UBA1^WT^ and UBA1^M41V^ THP-1 derived macrophages. Forty cells were quantified for n=6 independent experiments. SEM, Standard Error of the Mean; TLR, Toll-Like receptor; VEXAS, Vacuoles, E1 enzyme, X-linked, Autoinflammatory, Somatic. *P□<□0.05; **P□<□0.01; ***P□<□0.001, ****P□<□0.0001.

Similarly, UBA1^M41V^-derived macrophages exhibited similar phagocytic capacity compared to wild-type controls using pHrodo-labeled *E. coli* particles (**Figure 6C, Supplementary** Figure 6B), indicative of a functional endocytic machinery, which was further supported by preserved expression of the early and late endosomal markers Rab5 and Rab7 (**Figure 6D**). Intriguingly, we observed a decrease in EEA1 expression of the early endosome marker EEA1 (**Figure 6D)**. Decrease in EEA1 expression in inflammatory cells death has been reported ^33^ and seems to result from its secretion rather than transcriptional regulation.

Despite intact phagocytic uptake, UBA1^M41V^-derived macrophages displayed signs of lysosomal dysfunction. Specifically, we observed reduced expression of LAMP1 and LAMP2, as well as defective maturation of lysosomal proteases including cathepsins D, S, and B in UBA1^M41V^ macrophages (**Figure 6E**), pointing to impaired degradative processing. Immunofluorescence confirmed reduced LAMP1 signal concentrating on phagosomes containing engulfed apoptotic material (**Figure 6F and 6G**), reinforcing the conclusion of defective lysosomal fusion and resolution.

Collectively, these findings indicate that *UBA1*-mutated macrophages exhibit a pro-inflammatory phenotype and impaired efferocytosis, which could contribute to the chronic inflammation observed in VEXAS by hindering the clearance of dying cells and sustaining DAMP release in inflamed tissues ^34^.

## Discussion

Recent advances in the pathophysiology of VEXAS syndrome highlight its complexity. Ubiquitination, a key post-translational regulatory process, is involved in multiple cellular pathways ^14,15,35,36^ and defective ubiquitination in VEXAS syndrome affects various immune cell lineages across compartments, including HSPCs ^21^, circulating cells ^11,16,18^, and cells that infiltrate inflamed tissues ^12,13,37^. Our study identifies inflammatory cell death as a potential driver in the pathogenesis of VEXAS syndrome and reveals the specific contribution of *UBA1* mutations to this process. Using both patient samples and a genetically engineered monocytic cell model, we demonstrate that monocytes harboring the canonical UBA1^M41V^ mutation exhibit increased sensitivity to TNF-α-induced cell death and attenuated NF-κB responses upon TNF-α challenge. Our findings are supported by evidence of RIPK1 and MLKL phosphorylation in both patients’ primary monocytes and THP-1 mutated cells, as well as by the release of DAMPs such as HMGB1 in the context of UBA1 inhibition, which could propagate paracrine stress signals to neighboring wild-type cells, further impair hematopoiesis, and sustain inflammatory dominance. Importantly, we extend these mechanistic insights to the patient level, showing that whole blood from individuals with VEXAS exhibits increased spontaneous cytokine secretion, which was reversed by the RIPK1 inhibitor, necrostatin-1s. These observations are further supported by Adachi *et al.,* who demonstrated elevated cell death in monocytes from VEXAS patients, increased serum levels of HMGB1 and enhanced extracellular ATP secretion by monocytes, pointing toward the activation of inflammatory cell death in VEXAS ^17^. Furthermore, monogenic diseases involving impaired ubiquitination, such as LUBAC deficiencies, are characterized by reduced canonical NF-κB responses and an increased susceptibility to cell death mediated by TNF-α ^38,39^. These diseases exhibit a phenotype marked by severe systemic inflammation and recurrent infections, similar to that observed in VEXAS syndrome. These observations support a model in which *UBA1* mutations sensitize monocytes to inflammatory triggers by impairing key survival pathways and promoting cell death signaling. Given that RIPK1 inhibitors are currently in clinical development for other inflammatory diseases, our data provide a rationale for exploring this therapeutic avenue in VEXAS ^40^.

In parallel with enhanced inflammatory cell death, our data reveal that *UBA1* mutations disrupt the secretion of key pro-inflammatory cytokines following TLR stimulation. This defect was observed both ex vivo after stimulating patient whole blood and in vitro in *UBA1*-mutated THP-1 cells, and has been reported in CD14^+^ cells from VEXAS ^18^. Here, we demonstrate that this defect is not due to reduced TLR expression but rather to an impaired NF-κB response, indicating a signaling blockade downstream of TLR. Given the central role of ubiquitination in modulating the assembly and activation of TLR signaling components, including TRAF3, TRAF6 and RIPK1 ^14,31^, it is likely that UBA1 dysfunction impairs these processes. Overall, an impaired NF-κB-mediated response upon TNF-α and TLRs may contribute to the susceptibility to opportunistic infections observed in VEXAS patients.

Our study reveals that, beyond monocytes, *UBA1*-mutated macrophages exhibit profound functional abnormalities that likely contribute to tissue inflammation and disease persistence in VEXAS syndrome. THP-1-derived macrophages expressing the canonical *UBA1*^M41V^ mutation exhibited a heightened pro-inflammatory phenotype, characterized by increased secretion of proinflammatory cytokines. Transcriptomic profiling confirmed the enrichment of inflammatory gene signatures, including TNF-α/NF-κB signaling and type I/II interferon responses. Interestingly, upon polarization with LPS and IFN-γ, UBA1^M41V^ macrophages retained this exacerbated inflammatory profile but displayed reduced expression of key co-stimulatory molecules such as *CD80* and *CD86*, suggesting a state of hyperinflammation coupled with defective immune priming. In parallel, IL-4/IL-13-polarized UBA1^M41V^ macrophages exhibited impaired expression of immunomodulatory markers including *CD163, CD206, IL10*, and *MERTK*, as well as reduced secretion of IL-10. These findings indicate a polarization bias skewed toward inflammatory activation, with compromised anti-inflammatory or tissue-reparative responses. This imbalance may help explain the chronicity of inflammation in VEXAS tissues. Furthermore, the increased expression and secretion of chemokines such as CCL2, CCL3, and CCL20 by UBA1^M41V^ macrophages, mirroring patient plasma profiles, implicate these cells in promoting local recruitment of mutated myeloid cells. Together, these results suggest that *UBA1* mutations not only prime monocytes for inflammatory cell death but also reprogram macrophage responses in a way that reinforces inflammation, disrupts immune homeostasis, and limits resolution, features that may underlie the systemic and tissue-specific manifestations of VEXAS.

A critical finding of our study lies in the demonstration that UBA1^M41V^ macrophages, while preserving the capacity to engulf apoptotic bodies, exhibit a defect in the clearance and degradation of engulfed apoptotic material. Efferocytosis impairment has also been observed in sorted CD14□ cells from VEXAS patients ^18^. Our results suggest that defective expression of LAMP1 and LAMP2, as well as impaired maturation of cathepsins, leads to defective lysosomal fusion and function in *UBA1*-mutated macrophages. In the context of VEXAS, such defects have important immunological implications. Inefficient clearance of apoptotic debris results in secondary necrosis and substantial, non-resolving inflammatory responses ^34^, and defects in efferocytosis have been linked to various autoinflammatory disorders, particularly in the context of enhanced cell death ^34,41^. The histological features of VEXAS lesions, rich in nuclear debris, align with this pathophysiology and suggest that enhanced cell death and defective efferocytosis may contribute to tissue damage and the persistence of inflammatory signals together. In UBA1^M41V^ macrophages, the sustained production of inflammatory mediators, in the absence of an appropriate resolution response may contribute to the chronic and relapsing nature of tissue inflammation observed in VEXAS patients. These abnormalities could also explain why VEXAS patients are susceptible to intracellular pathogens, such as Mycobacterial and *Legionella Pneumoniae* infections, given the role of lysosomal function and trafficking in controlling such infections ^42–44^.

The study has several limitations. A previous THP-1 model harboring the same canonical *UBA1* mutation exhibited a distinct phenotype, characterized by the absence of inflammation, ubiquitination defects, and cell death. This cell line contains constitutive genomic editing from selected clones as compared to inducible pooled experiments used in this study. ^22^. Our model more closely resembles findings reported in the literature, particularly those derived from single-cell RNA sequencing data and analyses of sorted CD14□ from VEXAS patients suggests that constitutive editing may alter signaling pathways. Additionally, the limited number of patients and available samples, coupled with the monocytopenia characteristic of VEXAS syndrome, constrained the study of primary monocytes.

In conclusion, our study delineates a converging pathophysiological model in which *UBA1* mutations drive inflammation and tissue damage through dysregulated cell death, defective immune resolution, and altered intercellular communication. The identification of RIPK1 as a critical effector of both inflammatory cell death and DAMPs release highlights a promising therapeutic target. Finally, UBA1-mutant macrophages show lysosomal dysfunction and defective efferocytosis, potentially impairing inflammation resolution and susceptibility to intracellular pathogens.

## Material and methods

### Patients

Non-interventional study conducted between January 2021 and March 2024 within the RADIPEM biological sample collection framework. The collection and informed consent process were approved by the French Ministry of Research (No. 2019-3677) and the Cochin Hospital IRB (No. AAA-2021-08040). Written informed consent was obtained from all participants. The study followed Good Clinical Practice guidelines and the Declaration of Helsinki. For patients, genomic DNA was extracted from blood samples and the third exon of the gene was analyzed by a Sanger sequencing approach to detect the described mutations using specific primers as previously reported (Forward 5′TCCAAAGCCGGGTTCTAACT-3′/Reverse 5′-GGGTGTGCAGTAGGGAAA AA-3′) ^16^.

### Morphological analysis of skin biopsies of VEXAS patients

Sweet-syndrome-like skin VEXAS cases were identified from the archives of the Pathology department of Hôpital Cochin, Paris, France. Archived slides with hematoxylin, eosin, and saffron (HES) staining and immunohistochemistry (IHC) were reviewed by a dermatopathologist (P.S.).

### Immunofluorescence on THP-1 cells and CD14+ sorted cells

For THP-1 cells, cells were harvested at indicated time points and conditions by centrifugation at 300 × g for 5 minutes, then washed with 1xPBS. For CD14+ cells form patients, cells were sorted from 2mL fresh Whole Blood collected in EDTA tube and sorted using StraightFrom® Whole Blood CD14 MicroBeads and Whole Blood column kit (Milteny Biotec, #130-090-879) according to the manufacturer’s instructions. CD14+ sorted cells were then washed twice with 1xPBS. After counting THP-1 and CD14+ sorted cell suspension were adjusted to a final concentration of 1 × 10^6^ cells/mL. A volume of 100 µL of the cell suspension (approximately 1 × 10□ cells) was loaded into cytospin sample chambers (Cytospin funnels, Thermo Scientific) mounted onto microscope slides. Cytocentrifugation was performed at 2000 rpm (∼1000 × g) for 5 minutes at room temperature using a Cytospin 4 centrifuge (Thermo Scientific). After centrifugation, the slides were allowed to air dry for 10 minutes. Cells were then fixed with with 4% paraformaldehyde (PFA) for 10 minutes, then washed three times with phosphate-buffered saline (PBS) for 5 minutes at room temperature. Thereafter, cells were blocked for 1 hour at room temperature with a blocking solution consisting of PBS+3% (w/v) bovine serum albumin (BSA, Sigma-Aldrich) +0.1% Saponin (Sigma, #S7900). Subsequently, cells were incubated overnight at 4°C with primary antibodies in the blocking solution. Primary antibodies included anti phMLKL (1:200, Abcam, ab187091). The next day, cells were washed three times for 5 minutes with the blocking solution at room temperature. Donkey anti-rabbit Alexa 647 (1:500, Thermofisher Scientific) were subsequently incubated for 2 hours at RT in the dark. Then, cells were washed three times for 5 minutes with the blocking solution, and cell nuclei were stained with 1 *µ*g/ml Hoechst 33342 (Thermofisher Scientific, H3570) for 10 minutes in the dark. Afterward, fluorescent images were acquired using a Zeiss LSM900 inverted confocal laser scanning microscope with Airyscan 2 with a Plan-Apochromat 40×/1.2 water long-distance objective (Carl Zeiss Microscopy, Jena, Germany). The experiment was performed in sextuplicate, and at least 4 images were captured per experiment. The percentage of cell surface area positive for phMLKL was calculated using semiautomatic macros in ImageJ (version 1.53c, National Institutes of Health).

### Multiplex immunofluorescence of skin biopsy samples

Multiplex immunofluorescence was performed on 4□µm skin biopsy sections from VEXAS patient using the COMET™ platform (Lunaphore). Following deparaffinization, rehydration, and antigen retrieval, 28-plex Immunofluorescence stainings were performed using the COMET platform (Lunaphore Technologies SA, Switzerland). Images were acquired at 20× magnification and exported as stacked OME.tif files after automated background subtraction. The full experimental protocol and antibody references are detailed in the **Supplemental Materials and Supplemental Table 4.**

### RNA-Seq analysis of skin biopsies of VEXAS patients and healthy controls

Re-analysis of the publicly available dataset GSE245639 available at Gene Expression Omnibus, containing skin biopsy samples from VEXAS syndrome patients and healthy control patients was performed using R version 4.5.1 (R: A Language and Environment for Statistical Computing. R Foundation for Statistical Computing, Vienna, Austria. https://www.R-project.org). For complete protocol, see **Supplemental material**.

### Construction of a THP-1 VEXAS model cell line

To generate an inducible UBA1^KO^ cell line, human THP-1 cells were first infected with lentivirus packaged with FUCas9Cherry (Addgene, 70182) to constitutively express Cas9 and mCherry. Next, ten UBA1 targeting guide sequences were generated using CHOPCHOP software ^45^. The doxycycline-inducible FgH1tUTG (Addgene, 70183) plasmid was digested with BsmBI-V2 (NEB, R0739S), DNA gel electrophoresis purified, and T4 DNA Ligase (NEB, M0202S) ligated with the annealed product of a forward (5’-<u>TCCC</u>TAACCAGGACAACCCCGGTG-3’) and reverse (5’-<u>AAAC</u>CACCGGGGTTGTCCTGGTTA-3’) oligo containing BsmBI overhangs (underlined) and the UBA1-targeting sequence for the gRNA. The noted UBA1 target sequence was the most efficient in inducing KO of UBA1. This new plasmid, FgH1tUTG-UBA1, was then lentivirally packaged and infected into the Cas9 expressing THP-1. These cells were then sorted using FACS on a Sony SH800 Cell Sorter with a 130 μm sorting chip, selecting for dual positive mCherry (FUCas9Cherry) and GFP (FgH1tUTG-UBA1) THP-1 populations. Double positive cells were then induced with 2 μg/mL doxycycline with UBA1^KO^ validated by loss of endogenous UBA1a/b on immunoblot and eventual cell death.

Next, pHAGE_puro (Addgene, 118692) was modified to contain an IRES regulating puromycin expression and encode either a modified UBA1^WT^ (pHAGE-UBA1_WT-IRES-Puro) or UBA1^M41V^ (pHAGE-UBA1_M41V-IRES-Puro) cDNA product with a FLAG-HA tag. To avoid Cas9-mediated KO of the UBA1 transgene, the cDNA regions corresponding to the gRNA target sequence were modified to contain synonymous mutations (in bold) in the codons corresponding to amino acids 213-222 (5’-GT**C** AC**G** AA**A** GA**T** AA**T** CC**T** GG**A** GT**T** GT**G** AC**T**-3’). These constructs were then packaged into lentivirus, transduced into the dox-inducible UBA1^KO^ THP-1 cell line, and selected for 7 days in 1 μg/mL puromycin. After selection, trans-UBA1 expression was validated with immunoblot against HA-tag.

Finally, 2×10^6^ control UBA1^WT^ or UBA1^M41V^ expressing inducible UBA1^KO^ THP-1 cells were plated in a 10-cm diameter tissue culture dish with 12.5 mL of media containing 2 μg/mL doxycycline and grown for 6 days. For monocyte experimentation on UBA1^M41V^ and control UBA1^WT^, on day 6, cells were then split 1:3 into new media containing 2 μg/mL doxycycline and harvested on day 10. For monocyte experimentation on UBA1^KO^ and control UBA1^WT^cells were harvested on day 6. For macrophage experimentation on UBA1^M41V^ and control UBA1^WT^, induced cells were counted, spun at 300 xg for 10 mins, and supernatant removed. Cell pellets were then resuspended in new media containing 2 ug/mL doxycycline and 100ng/mL PMA at a concentration of 1×10^6^ cells/mL. These were then plated on tissue culture treated 6-well plates. Adherence and macrophage differentiation was observed by day 9 and cultures harvested on day 10.

### Analysis of cell vacuolization

Approximately 50,000 cells from the THP1 VEXAS lines were cytospin onto for 5□minutes at 2000□rpm with medium acceleration. Slides were dried 24h prior to staining with May-Grünwald Giemsa stain solution. Once dry, cells were cover slipped and images were acquired with a 40x oil objective on an upright, motorized Nikon Eclipse E600 microscope. These images were then quantified utilizing the ImageJ software and vacuolated cells were manually counted.

### RNA-seq analysis on THP-1 and THP-1 derived macrophages

THP-1 cells and THP-1 derived macrophages were collected at day 10 post Doxycyclin induction, washed in 1x PBS and RNA was extracted using Trizol (Thermofisher Scientific, 15596018). After RNA quantification and quality check, the libraries were prepared following the Stranded mRNA Prep protocol from Illumina, starting from 400 ng of high-quality total RNA. Paired end (2 × 59 bp) sequencing was performed on an Illumina Nextseq 2000 platform. For complete protocol on RNA seq analysis, see **supplemental material.**

Gene set enrichment analysis was performed as previously described ^46^. For specific analysis, the necroptosis gene list was obtained from^47^ and the NF-κB target gene list was adapted from^48^.

### Cell viability assay

THP-1 cells were seeded in 96-well plates and incubated with combinations of the following reagents: TNFα (50 ng/mL), the IAP inhibitor BV-6 (5 μM), the pan-caspase inhibitor Z-VAD-FMK (20 μM), the RIPK1 kinase inhibitor necrostatin-1 (20 μM), and the MLKL inhibitor necrosulfonamide (1 μM). After 24 hours, cell death was assessed using Zombie UV dye (Biolegend # 423108), and readout was performed with the Celigo Image Cytometer. Each condition was tested in triplicate.

### Annexin V apoptosis detection

THP-1 cells were incubated with TNF-α (50ng/mL) or left untreated. After 24 hours, supernatant was discarded, and cells were washed and stained using Annexin V and Propidium Iodide (PI) detection kit (Biolegend #640932) according to manufacturer instructions. Samples were transferred to flow cytometry tubes and analyzed using a BD LSFRFortessa-X20 (BD Biosciences) flow cytometer and data were analyzed with FlowJo Softwere (TreeStar, Inc.).

### HMGB1 reporter THP-1 cell line

THP-1 cells were cultured in a T-75 flask containing Roswell Park Memorial Institute 1640 Medium GlutaMAX™ Supplement medium (RPMI 1640 GlutaMAX™; Gibco) supplemented with 25□mM HEPES, 10% heat-inactivated fetal bovine serum (FBS), 100□U/mL penicillin, 100□μg/mL streptomycin, 100□µg/mL of Normocin (InvivoGen, ant-nr-05), and 100□µg/mL of Zeocin (InvivoGen, ant-zn-1p) according to the supplier’s instructions. For HMGB1 release measurement, THP-1 cells were cultured at 1×10^6^ cells/mL in 1□mL RPMI 1640 GlutaMAX supplemented medium. Cells were pre-treated with a selective inhibitor of ubiquitin-activating enzyme E1 PYR-41 (5□µM) with or without the pan-caspase inhibitor Z-VAD-FMK (20□μM), RIPK1 inhibitor necrostatin-1 (20□µM) or MLKL inhibitor necrosulfamid (1µM) or DMSO (control) for 1□h prior to incubation with recombinant human TNF-α (50□ng/ml). After 24□h, the luciferase activity was determined by measuring relative light units (RLUs) in a luminometer using QUANTI-Luc™ detection reagent. Each condition was tested in duplicate. For reactive reference, see **Supplementary Table 5.**

### Western blot analysis

Cells were stimulated with various reagents: TNF-α (50 ng/ml), LPS (100 ng/mL), R848 (2.5 µg/mL). The reactions were stopped with cold 1X PBS. THP-1 cells were washed with cold 1X PBS and lysed in a RIPA lysis buffer supplemented with a protease inhibitor (Roche) and phosphatase inhibitor (Roche). The isolated proteins were quantified using a Pierce BCA assay kit (ThermoFisher Scientific), subjected to SDS-PAGE, and transferred to a PVDF membrane. After blocking, the membranes were incubated with the primary antibodies (see **supplemental Table 6**. The results were visualized by chemiluminescence using species-specific HRP-linked secondary antibodies. Images of immunoblotting were analyzed with Fiji software.

### Cytokine and chemokines measurements

Supernatants from TruCulture and cell culture assays were analyzed for cytokines and chemokines with Luminex xMAP technology (R&D Systems) according to manufacturer recommendations as previously described ^16^.

### Whole blood stimulation assays

Whole blood stimulation assays with TLR agonists were performed as previously described ^32^. Briefly, whole blood was stimulated using TruCulture tubes preloaded with TLR agonists (poly(I:C), LPS, or R848), Necrostatin 1S (10µM) or unstimulated. After adding 1 mL of whole blood, tubes were incubated at 37°C for 22 hours. Stimulation was stopped and supernatants were collected and frozen at-80°C. See **Supplemental methods** for details.

### Cytometric Flux on THP-1 cells and THP-1 derived macrophages

For THP-1 cells, cells were harvested at indicated time points and conditions by centrifugation at 300 × g for 5 minutes, then washed in 1xPBS+2% BSA. For THP-1 cell surface staining of TLR 4, cells were resuspended in cell staining buffer (Biolegend, #420201) at a density of 1×10□ cells in 100 µL in flow cytometry tubes. Cells were incubated 30 min at 4°C in dark with anti TLR4 antibody (Biolegend, #312816), then washed twice in cell staining buffer and resuspended in 400µL before analysis. For THP-1 cells intracellular cell staining of TLR8, cells were fixed in IC Fixation buffer (Invivogen, #00-8222-49) 20 minutes at 4°C and washed twice in permeabilization buffer (Invivogen #00-8333-56) Cells were then resuspended in permeabilization buffer at a density of 1×10□ cells in 100 µL in flow cytometry tubes and stained with anti TLR8 antibody (Biolegend, #395509) for 30 min at 4°C in dark. Cells were washed twice with permeabilization buffer and resuspended in 400µL before analysis.

For macrophages, at indicated time points and conditions, cells were washed with 1xPBS then detached using Cell Stripper, centrifuged at 3000 g for 3 minutes, and resuspended in PBS containing 2% FBS at a density of 1×10□ cells in 500 µL in flow cytometry tubes. Cells were stained with anti-Mertk antibody (Biolegend, #367614) for 30 min at 4°C in dark, then washed twice with PBS + 2% BSA and resuspended in PBS + BSA 2% in 400µL before analysis. Samples were transferred to flow cytometry tubes and analyzed using a BD LSFRFortessa-X20 (BD Biosciences) flow cytometer and data were analyzed with FlowJo Softwere (TreeStar, Inc.).

### NF-***κ***B reporter THP-1 cell line

THP-1-Blue-NF-κB (InvivoGen) cells were cultured in a T-75 flask containing RPMI 1640, 2 mM L-glutamine, 25 mM HEPES, 10% heat-inactivated fetal bovine serum, 100 μg/ml Normocin (InvivoGen, ant-nr-05), Pen-Strep (100 U/ml-100 μg/ml) according to the supplier’s instructions. For monitoring the NF-κB signal transduction, THP-1 cells were cultured at 1×10^6^ cells/mL in 1□mL RPMI 1640 GlutaMAX supplemented medium. Cells were pre-treated with a selective inhibitor of ubiquitin-activating enzyme E1 PYR-41 (0.5, 5□and 10 µM) with or without the pan-caspase inhibitor Z-VAD-FMK (20□μM) and RIPK1 inhibitor necrostatin-1 (20□µM) or DMSO (control) for 1□h prior to incubation with TLR agonists (LPS, R848 and Pam3CSK4). After 24□h, Secreted embryonic alkaline phosphatase was determined by measuring optical density (OD) at 620-655 nm in a microplate reader using the QUANTI-Blue™ solution (InvivoGen). For reactive reference, see **Supplementary Table 5.**

### Cell migration assay

THP1 cells were seeded into the upper chamber of a 96-Well Monocyte Cell Migration Chamber (Sigma #CBA098) in RPMI GMAX medium without FBS (0% FBS). The lower chamber contained 150□µL of either RPMI GMAX 0% FBS (negative control), RPMI GMAX 10% FBS (positive control), or various chemokines (IL-8, CCL-2, CCL-3) diluted in RPMI GMAX 0% FBS. After 3 hours of incubation at 37□°C in 5% CO□, cells that had migrated to the lower chamber were counted with the Celigo Image Cytometer. Each condition was tested in triplicate.

### RNA Isolation and real-time quantitative reverse transcriptase PCR (RT-qPCR)

RNA was isolated from macrophages at day 10 post Doxycyclin induction using Trizol (Thermofisher Scientific, 15596018). Subsequently, 500 ng of RNA was reverse-transcribed into cDNA using the PrimeScript RT Reagent Kit (Takara Bio, Cat. #RR047A) according to the manufacturer’s instructions. Quantitative expression levels were determined by quantitative PCR using SYBR Green (Appliedbiosystems, A25742) on a StepOnePlus™ Real-Time PCR System (Thermofisher, 4376600). Expression levels were normalized to glyceraldehyde-3-phosphate dehydrogenase (GAPDH) levels using the delta–delta CT method. Six independent experiments were performed. All samples were measured in duplicate. Sequences of gene-specific primers are in **supplemental Table 7.**

### Efferocytosis and phagocytosis assay

For efferocytosis assay, HeLa cells were cultured in 1x PBS for 7 days to generate >90% annexin-V□ apoptotic cells (ACs), then labeled with Zombie Near Infra-Red dye (Biolegend) for 30 min, rinsed twice, and resuspended at 1×10□ cells/mL in RPMI 1640 GlutaMAX™ + 25□mM HEPES + 10% FBS. THP-1–derived macrophages and ACs were co-incubated (1:1 effector/target ratio) at a concentration of 1×10□ cells/mL in 6-well plates, for 2 hours. For phagocytosis assay, 10 μg of fluorogenic *E. coli* particles (P35360, Invitrogen) were co cultured with THP-1–derived macrophages in same conditions. Macrophages were then washed twice with 1× PBS to remove unbound material and incubated in fresh medium. At indicated time points, macrophages were washed to remove dead cells and detached using Cell Stripper (Corning) and resuspended in PBS +2% FBS. Samples were transferred to flow cytometry tubes and analyzed using a BD LSFRFortessa-X20 (BD Biosciences) flow cytometer and data were analyzed with FlowJo Softwere (TreeStar, Inc.).

For LAMP-1 immunofluorescent staining, efferocytosis assays was performed in µ-Slide 18 Well plates (ibidi, 81816) for 2 hours. After 2 washes, cells were incubated in fresh medium. At indicated time point, µ-Slide were washed, fixed with 4% PFA, permeabilized (PBS + 0.1% Triton), blocked (Tris-Buffered Saline with Tween 20 + FBS 3%), and stained with anti– LAMP1 antibody (1:100, Sigma, L1418) overnight at 4°C. Next day, slides were washed then stained with Donkey anti-rabbit Alexa 647 (1:500, Thermofisher Scientific) secondary antibody 2 hours. Fluorescent images were acquired on a Zeiss LSM900 confocal microscope with Airyscan 2 (40×/1.2 water objective). The experiment was performed in sextuplicate; ≥4 images per replicate were quantified using ImageJ macros to assess percentage of engulfed ACs area positive for LAMP1.

### Statistical analysis

All statistical analyses were performed using GraphPad Prism 9. As indicated in the Figures, results were analyzed in a two-tailed, unpaired or paired Student’s t test, a two-sided Mann– Whitney U test, or ANOVA (analysis of variance) tests followed by Dunn’s post-test for multiple group comparisons. For all analyses, the threshold for statistical significance was set to P < 0.05.

## Acknowledgments

We thank the following colleagues for their collaboration for their helpful discussion and diagnosis of VEXAS syndrome, all involved in the French VEXAS Study Group (FRENVEX), Sophie Georgin-Lavialle and Arsène Mekinian. We would also like to thank Hatem GABSI for his technique assistance.

## Author contributions

Experimental strategy and design: P.B, S.C, P-L.T, O.L, A.Z, O.K, D.B.B and B.T. Laboratory experiments: P.B, S.J.M, S.C, L.D, B.V, P.S, A.M, Q.D, F.P, N.R, K.C, M.P, B.T.

Statistical analyses: P.B., S.C and B.T. Manuscript writing: P.B. and B.T.

All authors critically revised the manuscript for important intellectual content and gave final approval for the version to be published. All authors agree to be accountable for all aspects of the work in ensuring that questions related to the accuracy or integrity of any part of the work are appropriately investigated and resolved.

## Data sharing statement

All data are available in the main text or the supplementary materials.

