## Supplemental Material and Method for "UBA1 Mutations Drive RIPK1-Mediated Cell Death and Monocyte Dysfunction in VEXAS Syndrome"

**Supplementary materials and methods**

*Immunohistochemistry of skin biopsy samples*

*RNA-Seq analysis of skin biopsies of VEXAS patients*

*Multiplex immunofluorescence of skin biopsy samples*

*RNA-seq analysis on THP-1 and THP-1 derived macrophages*

*Whole blood stimulation assays*

**Supplementary Material and Methods**

***Immunohistochemistry of skin biopsy samples***

Immunohistochemistry was conducted on tissue sections using the following steps: deparaffinization and antigen retrieval with the Leica Bond protocol (Leica Biosystems) employing Bond Epitope Retrieval Solution 2 (EDTA, pH 9.0) for 20 minutes. Antibodies targeting Myeloperoxidase (1:5000, GA51161-2, Dako), CD68 (1:100, GA61361-2, Dako) and CD15 (1:50, GA06261-2, Dako) served as the primary antibodies and were detected using the Polymer Refine Detection kit (Leica Biosystems) on a Leica Bond Autostainer (Leica Biosystems).

***Multiplex immunofluorescence of skin biopsy samples***

VEXAS skin tissue sections of 4 µm were baked overnight at 37°C to soften the paraffin. Deparaffinization was manually carried on in 3 successive baths of Clearene (Leica Biosystems, ref. 3803600E) and for 2 minutes each. Rehydration was performed by immersion in a graded ethanol series (100%, 90%, 70%, 50%) and a final 2-minute distilled water bath. Slides were stored in Tris-Buffered Saline 1X (Merck, ref T5912-1L) prior to antigen retrieval. Antigen retrieval was performed on the Epredia PT Module with the pH9 Dewax and HIER Buffer (Epredia™ TA-999-DHBH). Slides were dunked in this solution after a pre-warm cycle at 85°C. The antigen retrieval cycle was performed at 100°C for 1 hour, followed by a cooldown to 65°C.

28-plex Immunofluorescence stainings were performed using the COMET platform (Lunaphore Technologies SA, Switzerland). The first cycle of autofluorescence preceded 28 cycles of staining, imaging and elution. All cycles were automatically performed by the COMET platform once all the reagents were loaded. Each cycle was associated with 1 rabbit and 1 mouse primary antibodies that were both detected simultaneously with an anti-rabbit 647+ and an anti-mouse 555+ Alexa Fluor conjugated secondary antibody, respectively. After each staining cycle, elution of primary and secondary antibodies was performed with the provided Elution Buffer (Lunaphore, BU07-L). All antibody references are **presented in supplemental Table 4.**

For each sample, the image was taken at 20x magnification by the embarked microscope and its lasers (DAPI, Cy5, TRITC). A fully stacked OME.tif file was generated automatically once all cycles were completed. The OME.tif image was exported after we applied an automatic background subtraction algorithm on the Lunaphore COMET Viewer software.

***RNA-Seq analysis of skin biopsies of VEXAS patients and healthy controls***

Re-analysis of the publicly available dataset GSE245639 available at Gene Expression Omnibus, containing skin biopsy samples from VEXAS syndrome patients and healthy control patients was performed using R version 4.5.1 (R: A Language and Environment for Statistical Computing. R Foundation for Statistical Computing, Vienna, Austria. https://www.R-project.org).

Differential expression on the read counts were performed with the DESeq2 package version 1.48.1 to determine the differentially expressed genes between the two conditions. GSEA (Gene Set Enrichment Analysis) was performed using the fgsea package version 1.34.0 to determine whether gene sets from the Molecular Signatures Database, related to Hallmarks (v2025.1), C2 Canonical pathways (v2025.1) and C5 Gene Ontology (GO) (v2025.1), were randomly distributed throughout our ranked gene list or if they were located at the top or bottom of it. Genes were ranked according to the Wald test statistic of the differential expression. The fgseaMultilevel() function was used with the default parameters except minSize=15 and maxSize=500.

***RNA-seq analysis on THP-1 and THP-1 derived macrophages***

After RNA extraction, RNA concentrations were obtained using nanodrop or a fluorometric Qubit RNA assay (Life Technologies, Grand Island, New York, USA). The quality of the RNA (RNA integrity number) was determined on the Agilent 4200 TapeStation (Agilent Technologies, Palo Alto, CA, USA) as per the manufacturer’s instructions. To construct the libraries, 400 ng of high-quality total RNA sample (RIN >9) was processed using Stranded mRNA Prep kit (Illumina) according to manufacturer instructions. Briefly, after purification of poly-A containing mRNA molecules, mRNA molecules are fragmented and reverse- transcribed using random primers. Replacement of dTTP by dUTP during the second strand synthesis will permit to achieve the strand specificity. Addition of a single A base to the cDNA is followed by ligation of Illumina adapters. Libraries were quantified by Qubit (Invitrogen) and profiles were assessed using the DNA 1000 kit on an Agilent TapeStation 4200. Libraries were sequenced on an Illumina Nextseq 2000 instrument using 59 base-length reads in a paired-end mode. After sequencing, a primary analysis based on AOZAN software (ENS, Paris)*(49)* was applied to demultiplex and control the quality of the raw data (based of FastQC modules / version 0.11.5).

***Whole blood stimulation assays***

TruCulture tubes were prepared in batch with the indicated stimulus, resuspended in a volume of 1 mL media and stored at -20°C until use. Within one hour of collection, 1 mL of whole blood was added to each of the pre-warmed TruCulture tubes with or without following agents: TLR-3 agonist (Polyinosinic-polycytidylic acid (poly(I:C)), TLR-4 agonist (lipopolysaccharides (LPS)) and TLR-7/8 agonist (Resquimod (R848)), Necrostatin 1S (10µM) or unstimulated. TruCulture tubes were then placed in a dry block incubator and maintained at 37°C (±1°C) room air for 22 hours. At the end of the incubation period, the tubes were opened and a valve was inserted to separate the sedimented cells from the supernatant and to stop the stimulation reaction. Liquid supernatants were aliquoted and immediately frozen at -80°C until use.

**Supplemental Material References:**

1. S. Perrin, C. Firmo, S. Lemoine, S. Le Crom, L. Jourdren, Aozan: an automated post-sequencing data-processing pipeline. *Bioinformatics* **33**, 2212–2213 (2017).
