## Supplemental Figures legends for "UBA1 Mutations Drive RIPK1-Mediated Cell Death and Monocyte Dysfunction in VEXAS Syndrome"

**Supplementary figures legends:**

**Figure Supplemental S1. Skin lesions from VEXAS patients are characterized prominent infiltrate of CD68⁺ myeloid cells.**

Immunohistochemical staining of CD68, myeloperoxidase (MPO) and CD15 in the same skin biopsy. Illustrative picture is shown (Magnification x20, scale bar 200µm).

**Figure Supplemental S2. UBA1^M41V^ THP-1 cells model recapitulates VEXAS patient’s monocyte characteristics**

**(A**) Immunoblot analysis of UBA1 and total ubiquitin in the days following Doxycyclin induction of UBA1^WT^ and UBA1^KO^ THP-1 and (**B**) of UBA1^WT^ and UBA1^M41V^ THP-1 cells. (**C**) Representative images of vacuoles observed in UBA1^WT^ and UBA1^M41V^ THP-1 cells stained with May-Grünwald Giemsa in the day following Doxycyclin induction. (**D**) Quantification of number of vacuoles per cells of UBA1^WT^ and UBA1^M41V^ THP-1 cells. Mean ± SEM of n = 3 independent replicates (20 cells per replicates). Mann witney test was used to determine significance.

SEM, Standard Error of the Mean; VEXAS, Vacuoles, E1 enzyme, X-linked, Autoinflammatory, Somatic. *P < 0.05; **P < 0.01; ***P < 0.001, ****P < 0.0001.

**Figure Supplemental S3. Gasdermin D cleavage in UBA1 mutated models after TNFα challenge.**

(**A and C**) Representative immunoblot of Gasdermin D (GSDMD) and cleaved isoforms in UBA1^WT^ vs UBA1^M41V^(**A**) UBA1^WT^ vs UBA1^KO^ (**C**) THP-1 cells. Cells were treated with TNF-α (50 ng/ml) for 24H or left unstimulated. (**B and C**) Quantification of cleaved/ pro GSDMD in UBA1^WT^ vs UBA1^M41V^ (**B**) and UBA1^WT^ vs UBA1^KO^ (**C**) THP-1 cells. At least n=3 independent experiments are represented. 2 tailed p-values were determined with non-parametric ANOVA test followed by Dunn's post-test.

SEM, Standard Error of the Mean. *P < 0.05; **P < 0.01; ***P < 0.001, ****P < 0.0001.

**Supplementary Figure 4. UBA1^M41V^ THP-1-derived macrophages exhibit cell death and pro-inflammatory phenotype.**

**(A**) Immunoblot of UBA1a/b/c isoforms in UBA1^WT^ and UBA1^M41V^ THP-1 derived macrophages. Three independent experiments are represented. (**B**) Cytokine measurement in supernatant from UBA1^WT^ and UBA1^M41V^ THP-1 derived macrophages. Panels show 6 independent experiments (dots) and means ± SEM (histograms). (**C**) Representation of 15 Upregulated Hallmark pathways in UBA1^M41V^ THP-1 derived macrophages compared to UBA1^WT^ (n = 4 independently collected samples). Log (10) of q-value=0 was arbitrary set at 3. (**D**) GSEA analysis of Apoptosis (Hallmark) and Necroptosis (Culver-Cochran et al, Nature Communications, 2024) in UBA1^WT^ and UBA1^M41V^ derived macrophages. (**E**) Representative image of Celigo Image Cytometer Quantification of dead cell in UBA1^WT^ and UBA1^M41V^ THP-1 derived macrophages after zombie staining. GFP cells (circled in purple) are alive cells and Zombie cells (circled in yellow) are dead cells. (**F**) Quantification of dead cell percentages in UBA1^WT^ and UBA1^M41V^ THP-1 derived macrophages (*n=*6 independent experiments). (**G**) qRT-PCR analysis of pro inflammatory macrophage polarization markers and cytokine measurement in supernatant from UBA1^WT^ and UBA1^M41V^ THP-1 derived macrophages stimulated with IFN-y (20ng/ml) + LPS (1ng/ml). (**H**) qRT-PCR analysis of anti-inflammatory macrophage polarization markers and cytokine measurement in supernatant from UBA1^WT^ and UBA1^M41V^ THP-1 derived macrophages stimulated with IL-13 (20ng/ml) +IL-4 (20ng/ml) 48H. For (G) and (H), panels show 6 independent experiments (dots) and means ± SEM (histograms) and P values were determined by the Mann-Whitney test.

LPS, Lipopolysaccharides; SEM, Standard Error of the Mean; TLR, Toll-Like receptor; VEXAS, Vacuoles, E1 enzyme, X-linked, Autoinflammatory, Somatic; WT, Wild-Type. *P < 0.05; **P < 0.01; ***P < 0.001, ****P < 0.0001.

**Figure Supplemental S5. Plasma from VEXAS patients and supernatant from UBA1 ^M41V^ derived macrophage secrete more chemokines and UBA1^M41V^ THP-1 cells displayed increased migration in response to chemokines.**

(**A**) Chemokines level in plasma in patients with VEXAS (n=40) and healthy controls (n=10). Each dot represents a single patient. Panels show individual data (dots) and means ± SEM (histograms). Mann witney test was used to determine significance. (**B**) Chemokines level in supernatant from UBA1^WT^ and UBA1 ^M41V^ derived macrophage. Panels show n=6 independent experiments (dots) and means ± SEM (histograms). Mann witney test was used to determine significance. (**C**) UBA1^WT^ and UBA1^M41V^ THP-1 cells migration in presence of RPMI GMAX +0% FBS (negative control), RPMI GMAX +10% FBS (positive control) and tested condition; IL-8 (100ng/ml) CCL-2 (0.5µg/ml) and CCL-3 (50ng/mL). After 3H, cells that had migrated were counted with the Celigo Image Cytometer. Each condition was tested in triplicate. Panels show n=6 independent experiments (dots) and means ± SEM (histograms). Mann witney test was used to determine significance. (**D**) THP-1^WT^ cells migration in presence of supernatant of UBA1^WT^ and UBA1 ^M41V^ derived macrophage (48H culture). After 3H, cells that had migrated were counted with the Celigo Image Cytometer. Each condition was tested in triplicate. Panels show n=6 independent experiments (dots) and means ± SEM (histograms).

*P < 0.05; **P < 0.01; ***P < 0.001, ****P < 0.0001 SEM, Standard Error of the Mean

**Figure Supplemental S6. Gating strategy of flow cytometric analysis of efferocytosis and phagocytosis assays.**

(**A**) Representative images of the gating strategy used for flow cytometric analysis of apoptotic bodies clearance in UBA1^WT^ and UBA1^M41V^ THP-1 derived macrophages using HeLa dead cells stained with zombie dye as target (1:1 ratio with live Macrophages) showed in Figure 7. Repartition of dead Macrophages was determined using Zombie staining in Macrophages not cultured with HeLa dead cells. (**B**) Representative images of the gating strategy used for the flow cytometric analysis of the phagocytic capacity of UBA1^WT^ and UBA1^M41V^ THP-1 derived macrophages using pHrodo *E. coli* as target particles showed in Figure 6.
