## Supplemental Tables for "UBA1 Mutations Drive RIPK1-Mediated Cell Death and Monocyte Dysfunction in VEXAS Syndrome"

**Supplementary tables**

**Supplemental Table 1. Demographic and clinical characteristics of participants included in the analysis of white blood cell count.**

|  | **Healthy controls (n=14)** | **VEXAS**  **(n=52)** |
| --- | --- | --- |
| **Demographic characteristics** |  |  |
| Male, n (%) | 14 (100) | 52 (100) |
| Age at disease onset, median (IQR), years | - | 71.2 (65.4-78.5) |
| Age at sampling, median (IQR), years | 60.5 (59.6-73.8) | 74 (67.9-79.9) |
| **Somatic UBA1 variants** | 0 (0) | 52 (100) |
| p. Met41Thr (c.122T>C), n (%) | - | 24 (46.1) |
| p. Met41Leu (c.121A>C), n (%) | - | 10 (19.2) |
| p. Met41Val (c.121A>G), n (%) | - | 12 (23) |
| Splice motif mutation, n (%) | - | 6 (11.5) |
| **Key clinical features** |  |  |
| Fever, n (%) | - | 31 (59.6) |
| Skin involvement, n (%) | - | 41 (78.8) |
| Arthralgia/arthritis, n (%) | - | 32 (61.5) |
| Pulmonary infiltrate, n (%) | - | 15 (28.8) |
| Ear/nose chondritis, n (%) | - | 27 (51.9) |
| Venous thromboembolism, n (%) | - | 23 (44.2) |
| **Hematological conditions:** |  |  |
| Macrocytic anemia, n (%) | - | 44 (84.6) |
| Thrombopenia, n (%) | - | 23/50 (46) |
| Myelodysplastic syndrome, n (%) | - | 26 (50) |
| **Laboratory findings at time of sampling** |  |  |
| Hemoglobin, median (IQR), g/L | 15.7 (15.3-15.9) | 9.9 (8.7-10.9) |
| Mean corpuscular volume, median (IQR), fL | 90 (89-94) | 106 (99-113) |
| Leukocytes, median (IQR), x10^9^/L | 6.7 (6.3-7.7) | 4.1 (2.5-5.6) |
| Neutrophil, median (IQR), x10^9^/L | 4 (3.4-4.7) | 3 (1.8-4.4) |
| Lymphocytes, median (IQR), x10^9^/L | 2 (1.4-2.9) | 0.75 (0.45-1.2) |
| Monocytes, median (IQR), x10^9^/L | 0.5 (0.3-0.6.3) | 0.2 (0.14-0.34) |
| **Current treatment at time of sampling** |  |  |
| Glucocorticoids, n (%) | 0 (0) | 40/51 (78.4) |
| Prednisone dose, median (IQR), mg/day | 0 (0) | 10 (5-17.5) |
| Synthetic DMARDs, n (%) | 0 (0) | 8/51 (15.7) |
| Biologic or target synthetic DMARDs, n (%) | 0 (0) | 11/51 (21.6) |
| 5-azacytidine, n (%) | 0 (0) | 4/51 (7.8) |

**Supplemental Table 2. Main characteristics of participants included in the Whole Blood Stimulation assay.**

| **Patient ID** | **Gender** | **Age (years)** | ***UBA1* status** | **Treatment at sampling** | **Disease status** |
| --- | --- | --- | --- | --- | --- |
| Healthy control 1 | Male | 45 | WT | - | - |
| Healthy control 2 | Male | 45 | WT | - | - |
| Healthy control 3 | Female | 57 | WT | - | - |
| Healthy control 4 | Female | 50 | WT | - | - |
| Healthy control 5 | Female | 50 | WT | - | - |
| VEXAS 1 | Male | 74 | p. Met41Leu (c.121A>C) | 5-azacytidine | Inactive |
| VEXAS 2 | Male | 60 | p. Met41Thr (c.122T>C) | Prednisone 10 mg/d + Ruxolitinib | Active |
| VEXAS 3 | Male | 85 | p. Met41Leu (c.121A>C) | - | Active |
| VEXAS 4 | Male | 79 | p. Met41Thr (c.122T>C) | Prednisone 15 mg/d | Active |
| VEXAS 5 | Male | 62 | p. Met41Leu (c.121A>C) | 5-azacytidine | Active |
| VEXAS 6 | Male | 79 | p. Met41Leu (c.121A>C) | Prednisone 10 mg/d | Active |
| VEXAS 7 | Male | 80 | p. Met41Thr (c.122T>C) | - | Inactive |
| VEXAS, Vacuole, E1 enzyme, X-linked, autoinflammatory, somatic; UBA1, Ubiquitin Like Modifier Activating Enzyme 1; WT, Wild-Type. | | | | | |

**Supplemental Table 3. Demographic and clinical characteristics of participants included in the analysis of chemokines measurement.**

|  | **Healthy controls (n=10)** | **VEXAS**  **(n=40)** |
| --- | --- | --- |
| **Demographic characteristics** |  |  |
| Male, n (%) | 10 (100) | 40 (100) |
| Age at disease onset, median (IQR), years | - | 71.9 (66-78.7) |
| Age at sampling, median (IQR), years | 60.7 (55.6-78.1) | 74 (68.5-79.9) |
| **Somatic UBA1 variants** | 0 (0) | 40 (100) |
| p. Met41Thr (c.122T>C), n (%) | - | 18 (45) |
| p. Met41Leu (c.121A>C), n (%) | - | 6 (15) |
| p. Met41Val (c.121A>G), n (%) | - | 10 (25) |
| Splice motif mutation, n (%) | - | 6 (15) |
| **Key clinical features** |  |  |
| Fever, n (%) | - | 27 (67.5) |
| Skin involvement, n (%) | - | 31 (77.5) |
| Arthralgia/arthritis, n (%) | - | 26 (65) |
| Pulmonary infiltrate, n (%) | - | 14 (35) |
| Ear/nose chondritis, n (%) | - | 19 (47.5) |
| Venous thromboembolism, n (%) | - | 17 (42.5) |
| **Laboratory findings at time of sampling** |  |  |
| Hemoglobin, median (IQR), g/L | 15.6 (14.4-15.8) | 10 (8.6-10.7) |
| Leukocytes, median (IQR), x10^9^/L | 6.7 (4.7-6.9) | 3.9 (2.5-5.6) |
| **Current treatment at time of sampling** |  |  |
| Glucocorticoids, n (%) | 0 (0) | 34 (85) |
| Prednisone dose, median (IQR), mg/day | 0 (0) | 10 (7-19.4) |
| Synthetic DMARDs, n (%) | 0 (0) | 8 (20) |
| Biologic or target synthetic DMARDs, n (%) | 0 (0) | 9 (22.5) |
| 5-azacytidine, n (%) | 0 (0) | 4 (10) |

DMARDs, Disease-modifying antirheumatic drugs; IQR, interquartile range; VEXAS, Vacuole, E1 enzyme, X-linked, autoinflammatory, somatic; UBA1, Ubiquitin Like Modifier Activating Enzyme 1.

**Supplemental Table 4 Antibody used for the** **multiplex immunofluorescence of skin biopsy samples.**

| **Target** | **Reference** | **Company** |
| --- | --- | --- |
| CD45 | MR10090 | Lunaphore |
| CD3 | A0452 | Dako |
| CD4 | MR10020 | Lunaphore |
| CD8 | MR10030 | Lunaphore |
| CD20 | MR10050 | Lunaphore |
| CD56 | MR10060 | Lunaphore |
| CD68 | MR10080 | Lunaphore |
| CD163 | 163M-1 | Cell Marques |
| CD11c | MR10070 | Lunaphore |
| FOXP3 | MR10040 | Lunaphore |
| αSMA | MR10100 | Lunaphore |
| CD31 | ab9498 | Abcam |
| PDL1 | MR10130 | Lunaphore |
| PD1 | ab186928 | Abcam |
| MUM1 | M7259 | Dako |
| HLA-DR | sc-53319 | Santa Cruz |
| MLKL | MABC1636 | Merck |
| pRIPK1 | #96323, BSA & Azide Free | Cell Signaling |
| cCasp3 | #9664 | Cell Signaling |
| NLRP3 | HPA012878 | Merck |
| cGSDMD | #37349, BSA & Azide Free | Cell Signaling |
| TREM2 | #62264 BSA & Azide Free | Cell Signaling |
| FOLR2 | MA5-26933 | ThermoSci |
| TFRC | ab269513 | Abcam |
| GPX4 | ab125066 | Abcam |
| Myeloperoxidase (MPO) | A0398 | Dako |
| Cytokeratin | MR10140 | Lunaphore |
| CXCR5 | bsm-54319R | Bioss |

**Supplement Table 5. List of reagents.**

| **Reagents** | **Reference** | **Company** |
| --- | --- | --- |
| TNF-α | 210-TA | Bio-Techne |
| LPS | L2630 | Sigma |
| R848 | tlrl-r84 | Invivogen |
| Pam3CSK4 | tlrl-pms | Invivogen |
| CCL-2 | SRP 3109 | Sigma |
| CCL3 | 300-08 | Perprotech |
| IL-8 | SRP 30-98 | Sigma |
| IFN-y | 285-IF-100 | Bio-Techne |
| IL-13 | 213-ILB | Perprotech |
| IL-4 | BT-004 | Perprotech |
| Z-VAD-FMK | tlrl-vad | InvivoGen |
| PYR-41 | T6629 | TargetMo |
| Necrostatin-1 | inh-ncst1 | InvivoGen |
| Necrostatin- 1S | HY-14622A | MedChemExpress |
| Necrosulfamid | HY-100573 | MedChemExpress |
| BV6 | S7597 | Selleckchem |

**Supplemental Table 6. List of antibodies used in Western Blot.**

| **Target** | **Reference** | **Company** |
| --- | --- | --- |
| Caspase-3 | #9662 | Cell Signaling Technology |
| Cleaved caspase 3 | #9661 | Cell Signaling Technology |
| Caspase 8 | #9746 | Cell Signaling Technology |
| Cleaved caspase 8 | #9496 | Cell Signaling Technology |
| RIPK1 | #3493 | Cell Signaling Technology |
| Phospho-RIPK1 (Ser166) | #44590 | Cell Signaling Technology |
| Gasdermin D | #97558 | Cell Signaling Technology |
| Cleaved Gasdermin D | #36425 | Cell Signaling Technology |
| UBE1a/b/c | #4891 | Cell Signaling Technology |
| Ubiquitin (P4D1) | #3936 | Cell Signaling Technology |
| NF-κB p65 | sc-372 | Santa-Cruz |
| phospho-NF-κB p65 (Ser536) | #3033 | Cell Signaling Technology |
| IKβα | #4814 | Cell Signaling Technology |
| IKKα | #2682 | Cell Signaling Technology |
| IKKβ | #2684 | Cell Signaling Technology |
| Phospho IKKα/β (ser176/180) | #2694 | Cell Signaling Technology |
| LAMP1 | #3243 | Cell Signaling Technology |
| LAMP2 | #49067 | Cell Signaling Technology |
| Cathepsin S | #25084 | Cell Signaling Technology |
| Cathepsin D | #74089 | Cell Signaling Technology |
| Cathepsin B | #31718 | Cell Signaling Technology |
| Rab5 | #3547 | Cell Signaling Technology |
| Rab7 | #9367 | Cell Signaling Technology |
| EEIA | #3228 | Cell Signaling Technology |
| β-Actin | #4970 | Cell Signaling Technology |

**Supplement Table 7. List of primers used for RT qPCR assays.**

| **Primer Name** | **Sense** | **Sequence** |
| --- | --- | --- |
| iNOS | F | CTGGCAAGCCCAAGGTCTAT |
|  | R | TCCCCGCAAACATAGAGGTG |
| CXCL-10 | F | CCACGTGTTGAGATCATTGCTA |
|  | R | TTGATGGCCTTCGATTCTGG |
| CD80 | F | CTCTTGGTGCTGGCTGGTCTTT |
|  | R | GCCAGTAGATGCGAGTTTGTGC |
| CD86 | F | AGACGCGGCTTTTATCTTCACCTTT |
|  | R | CGAGGCCGCTTCTTCTTCTTCCAT |
| CD163 | F | GGCCCCTCGGAGGAGACCTGGATCACA |
|  | R | TCCCCCAGGAACCTCCATGCCAGATC |
| CD206 | F | GGGCCAAGCTTCTCTGGAATGTCT |
|  | R | AGTGGCTCAACCCGATATGACAGAA |
| MERTK | F | CAGCCCGAAGACTGCCTGGATGAA |
|  | R | GGGCGGTCTAAGGGATCGGTTCT |
| IL-10 | F | TTGCTGGAGGACTTTAAGGGTT |
|  | R | CACATGCGCCTTGATGTCTG |
| GAPDH | F | AATGCTCCCAGAAATGGCAC |
|  | R | CCTCCACTGAACCCTGGAAA |
