## Supplementary figures and images for "UBA1 Mutations Drive RIPK1-Mediated Cell Death and Monocyte Dysfunction in VEXAS Syndrome"

### Supplemental Figure 1

CD68

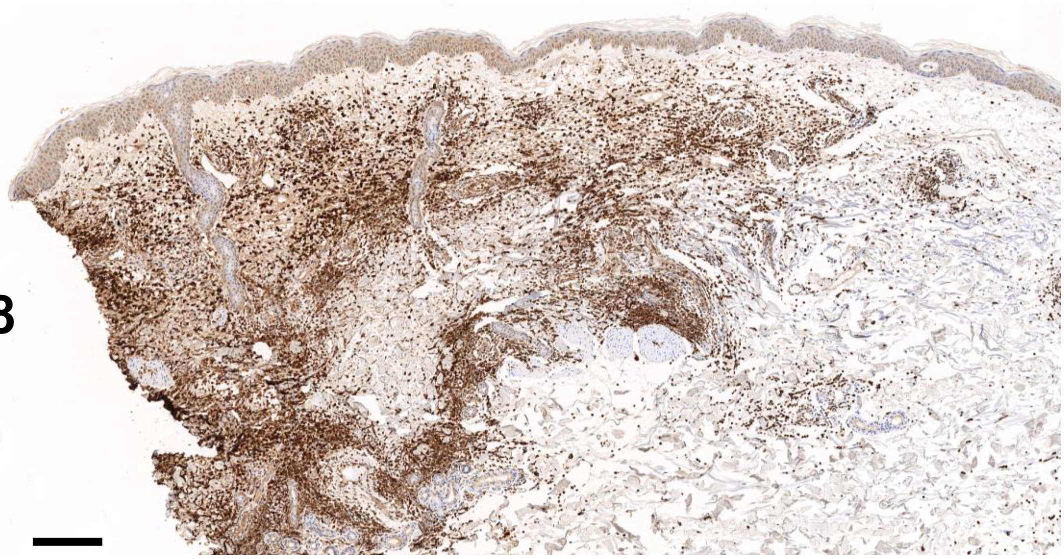

CD15

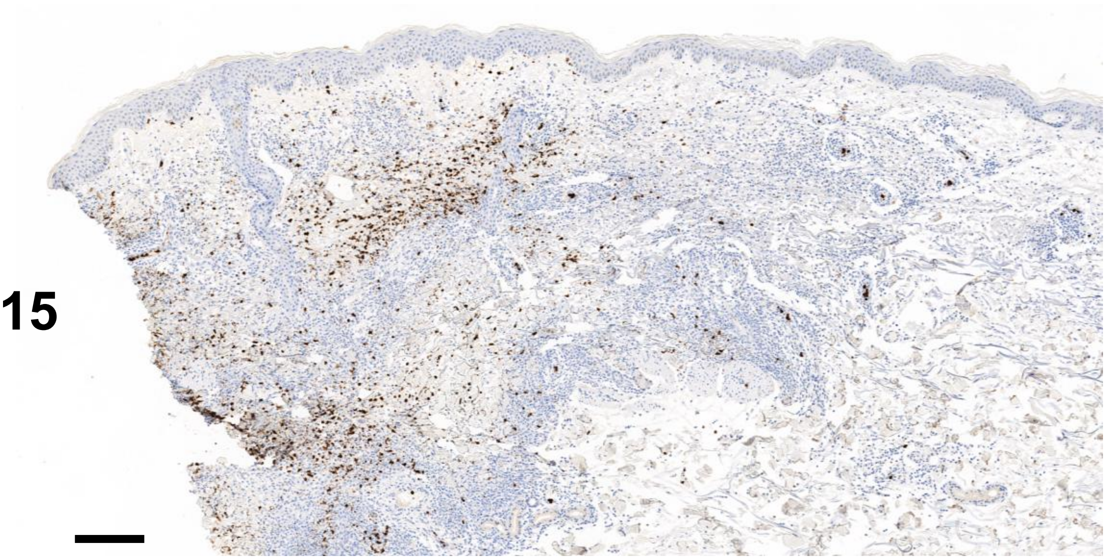

MPO

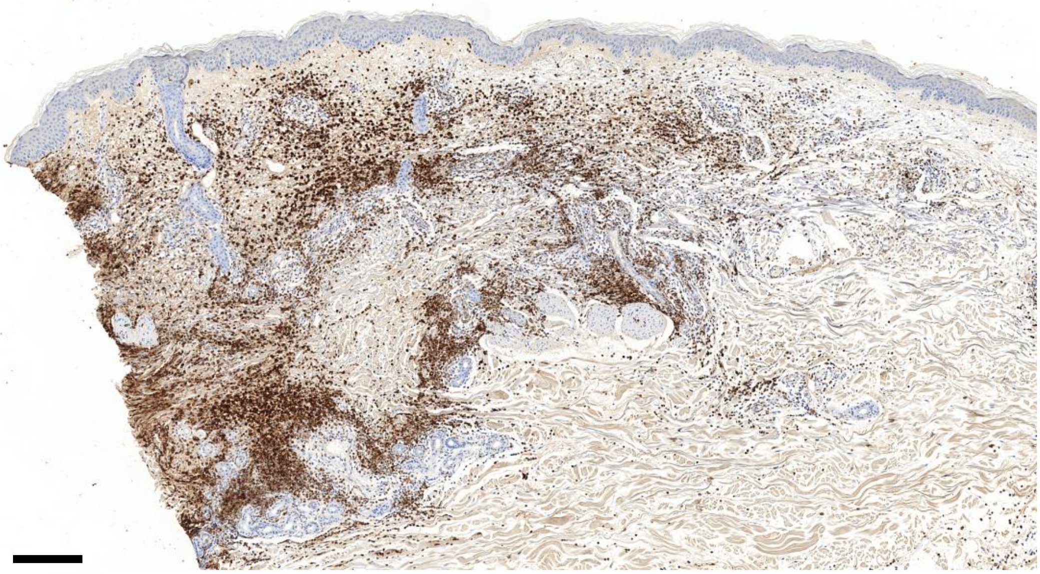

### Supplemental Figure 2

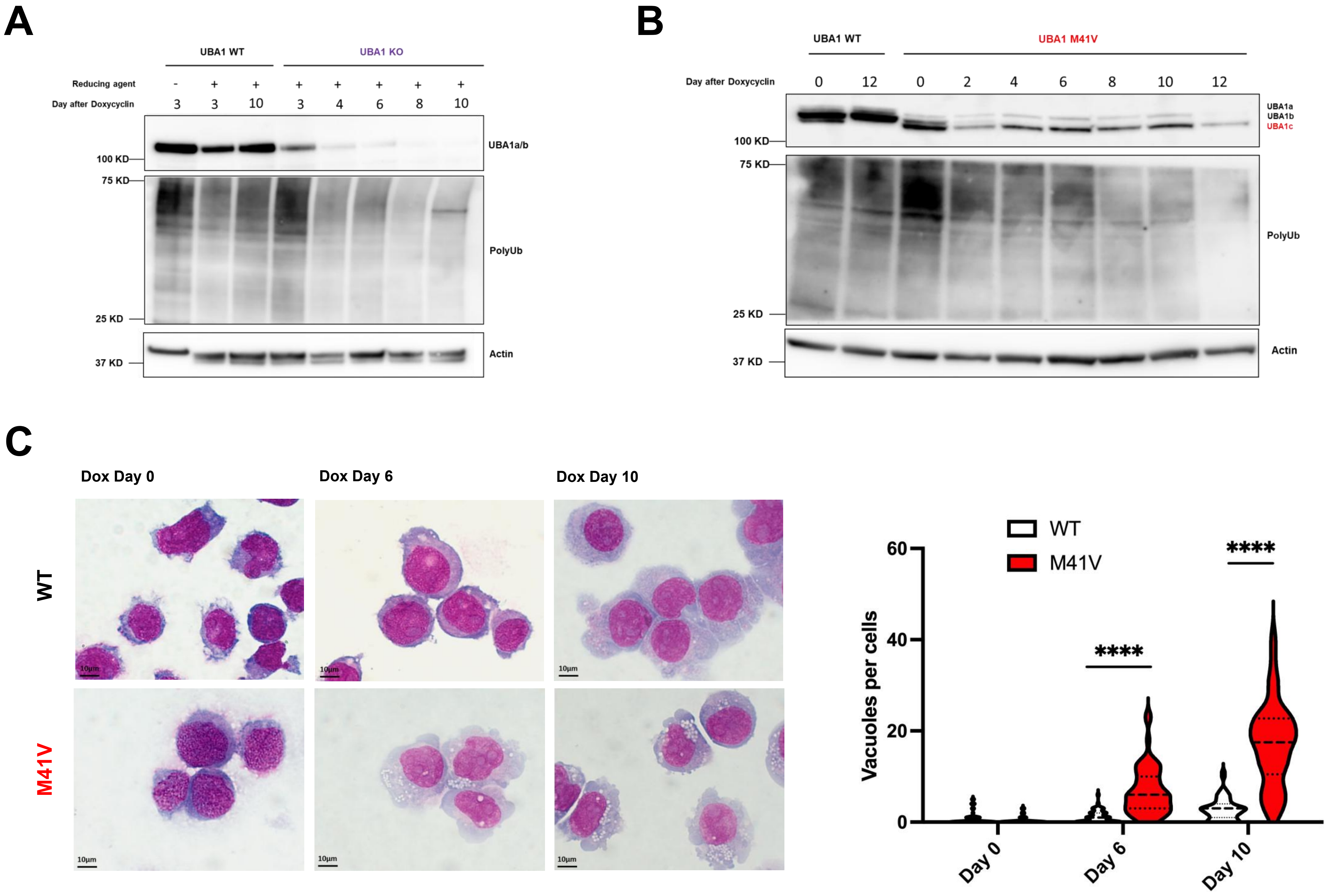

### Supplemental Figure 3

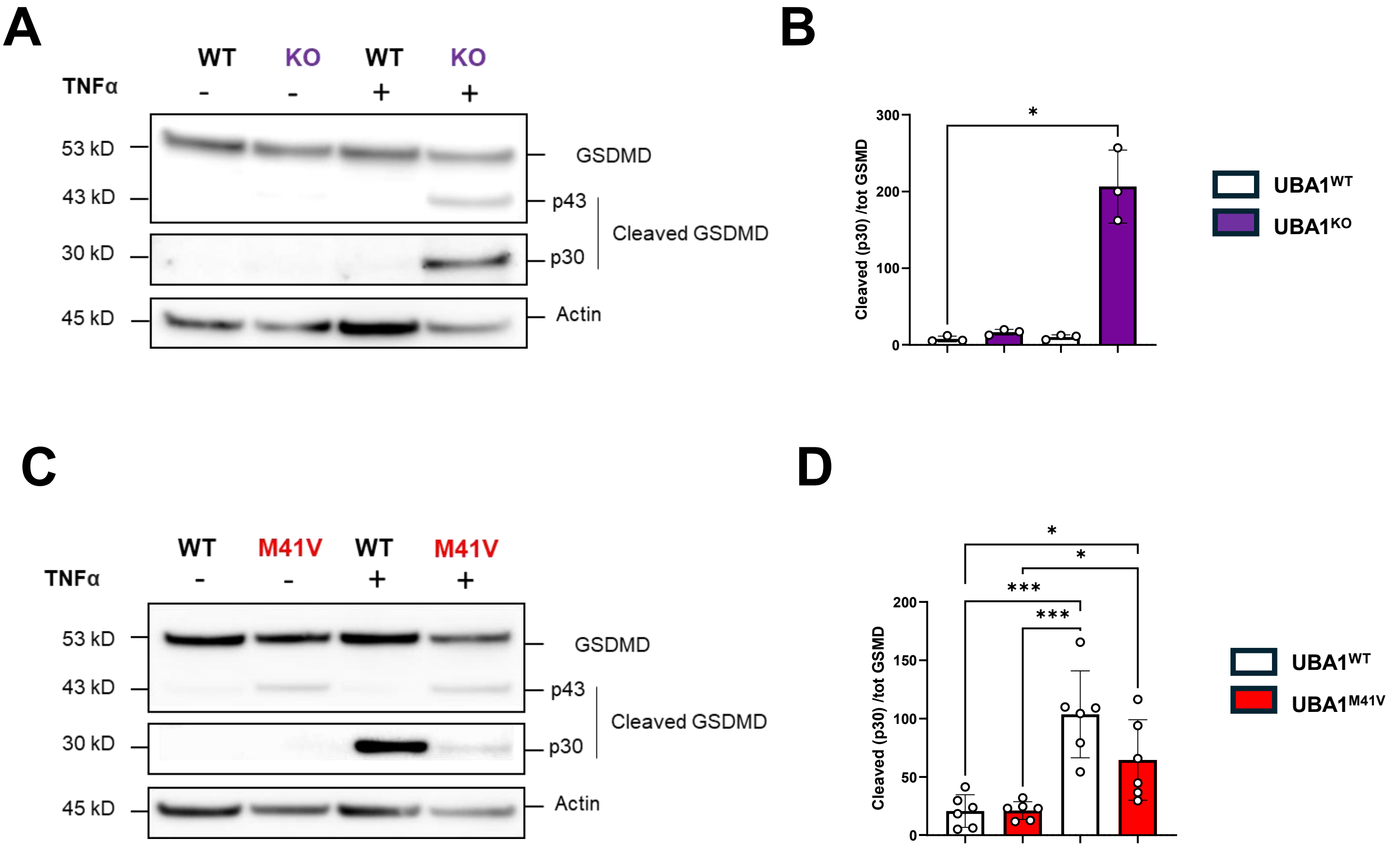

### Supplemental Figure 4

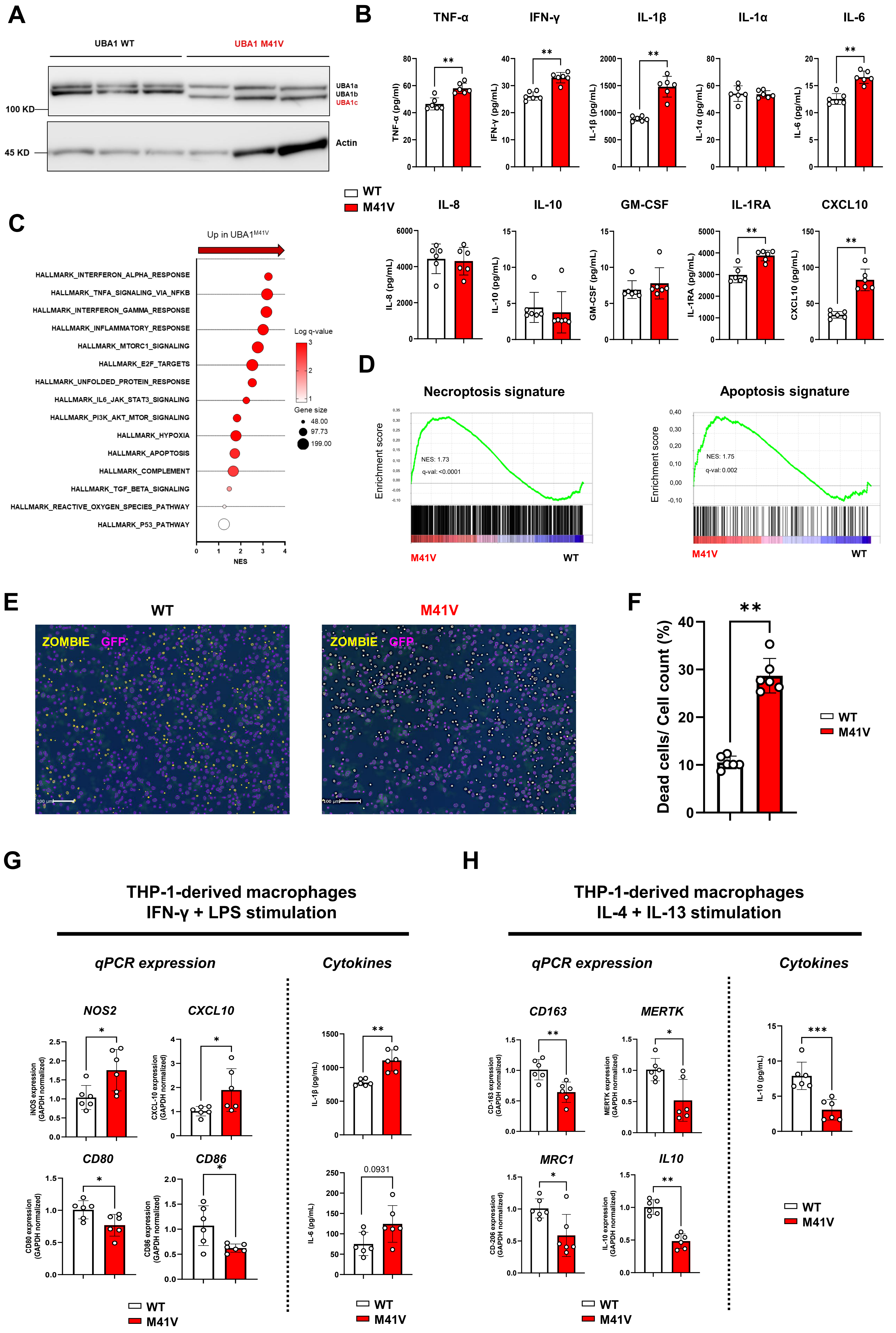

### Supplemental Figure 5

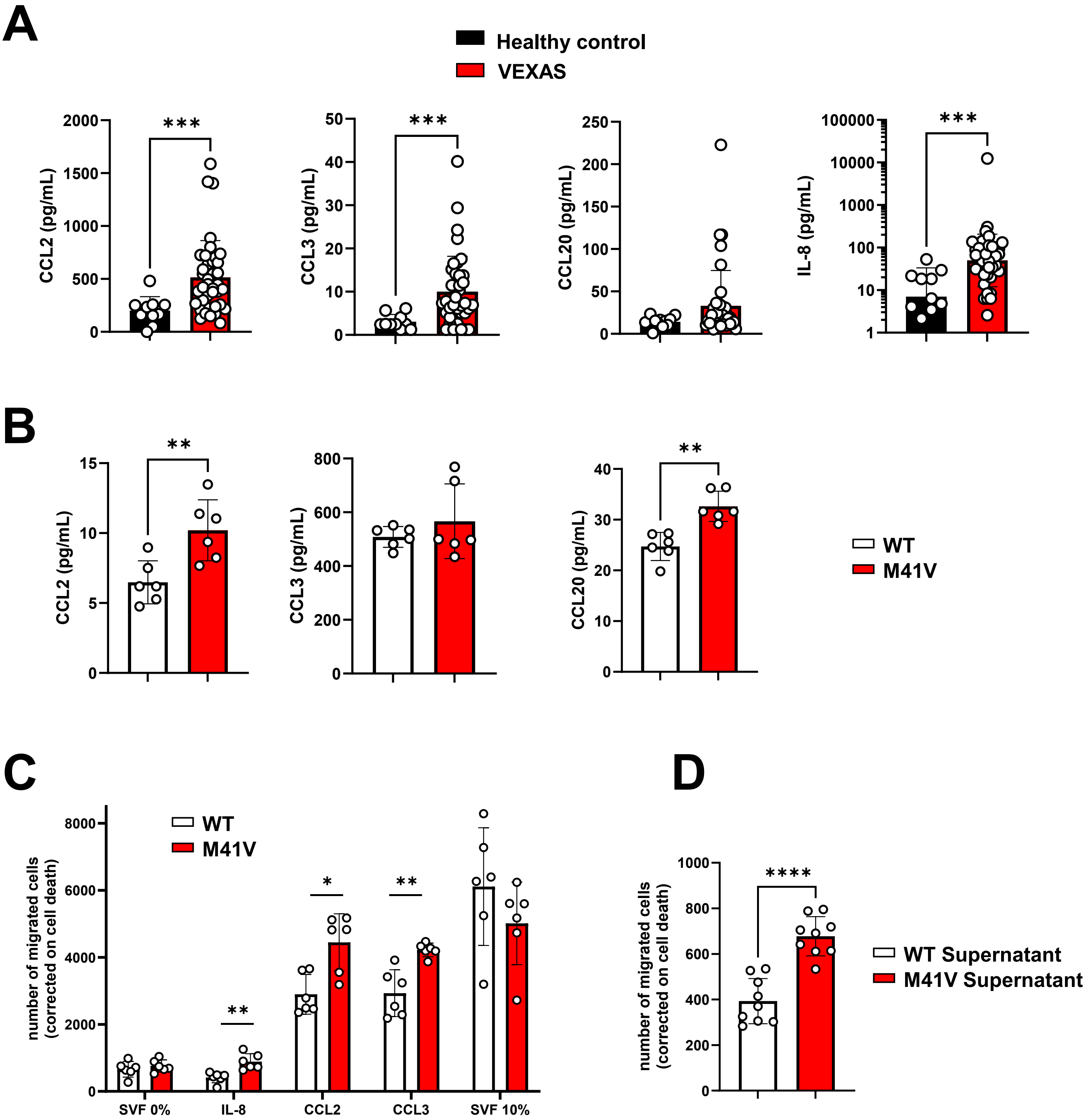

### Supplemental Figure 6

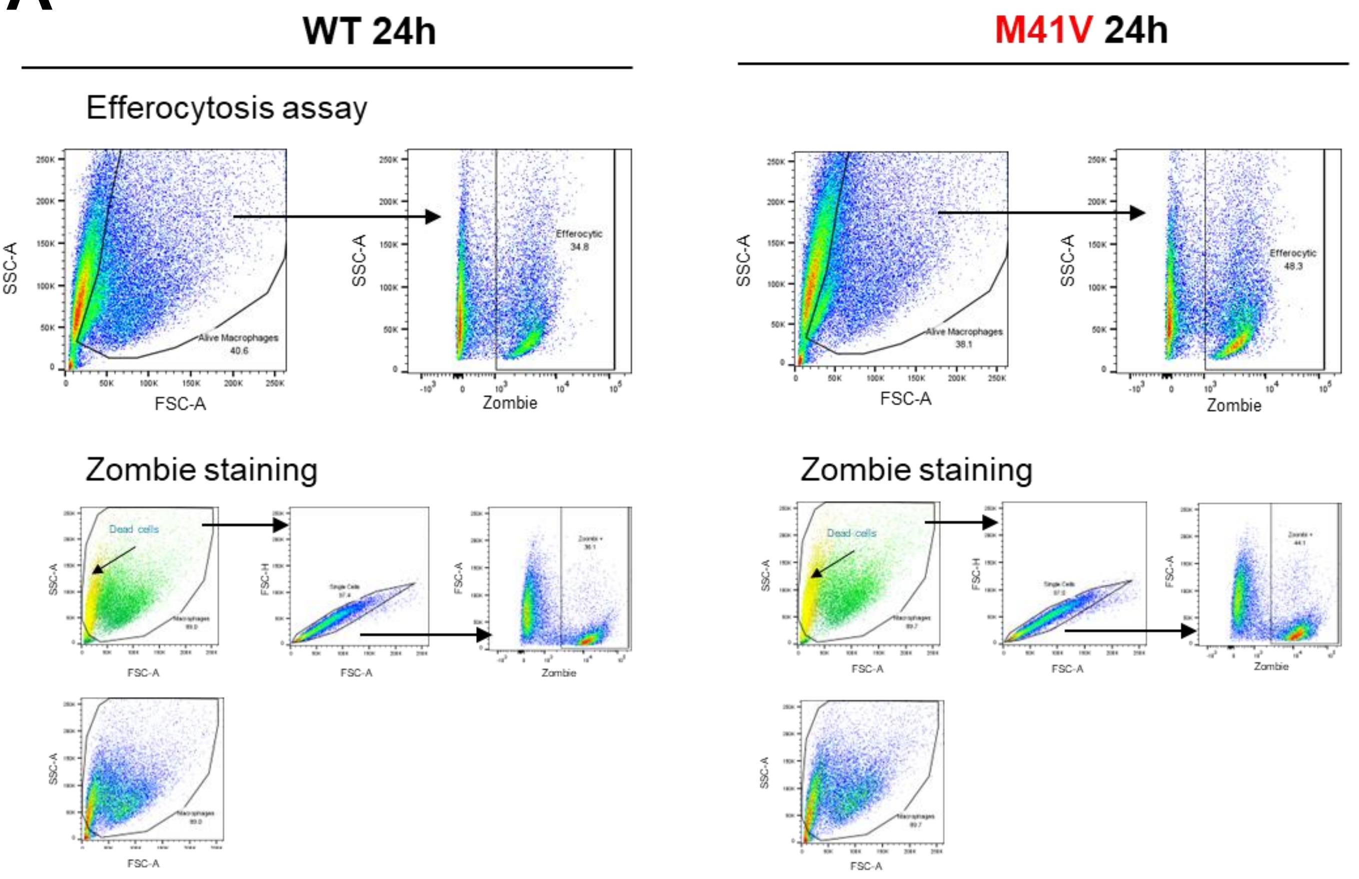

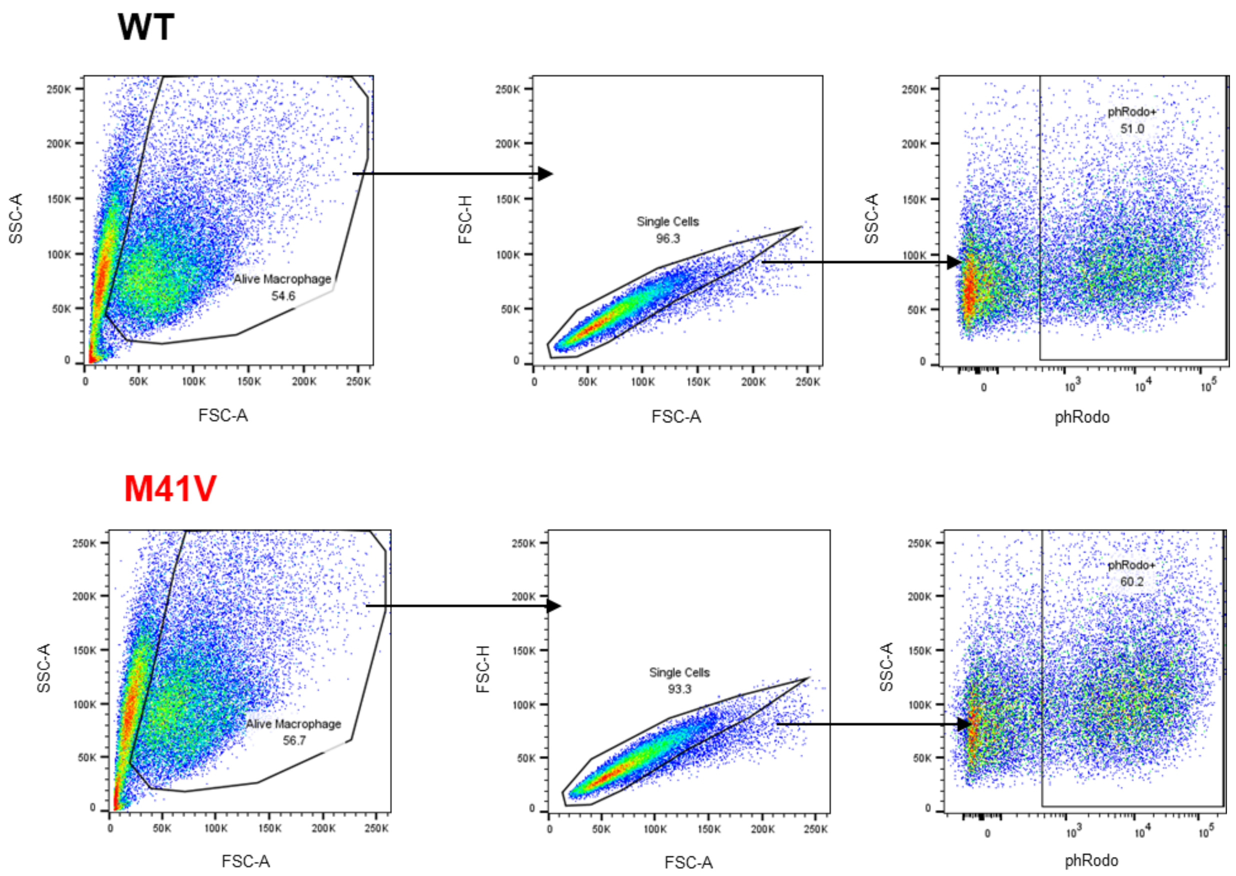
